# SUMO2 Inhibition Reverses Aberrant Epigenetic Rewiring Driven by Synovial Sarcoma Fusion Oncoproteins and Impairs Sarcomagenesis

**DOI:** 10.1101/2024.09.23.614593

**Authors:** Rema Iyer, Anagha Deshpande, Aditi Pedgaonkar, Pramod Akula Bala, Taehee Kim, Gerard L. Brien, Darren Finlay, Kristiina Vuori, Alice Soragni, Rabi Murad, Aniruddha J. Deshpande

## Abstract

Synovial Sarcoma (SySa) is an aggressive soft tissue sarcoma that accounts for 5 – 10% of all soft tissue sarcomas. Current treatment involves radiation and radical surgery including limb amputation, highlighting the urgent need to develop targeted therapies. We reasoned that transcriptional rewiring by the fusion protein SS18-SSX, the sole oncogenic driver in SySa, creates specific vulnerabilities that can be exploited for treatment. To uncover genes that are selectively essential for SySa, we mined The Cancer Dependency Map (DepMap) data to identify genes that specifically impact the fitness of SySa compared to other tumor cell lines. Targeted CRISPR library screening of SySa-selective candidates revealed that the small ubiquitin-like modifier 2 (SUMO2) was one of the strongest dependencies both *in vitro* as well as *in vivo*. TAK-981, a clinical-stage small molecule SUMO2 inhibitor potently inhibited growth and colony-forming ability. Strikingly, transcriptomic studies showed that pharmacological SUMO2 inhibition with TAK-981 treatment elicited a profound reversal of a gene expression program orchestrated by SS18-SSX fusions. Of note, genetic or pharmacological SUMO2 inhibition reduced global and chromatin levels of the SS18-SSX fusion protein with a concomitant reduction in histone 2A lysine 119 ubiquitination (H2AK119ub), an epigenetic mark that plays an important role in SySa pathogenesis. Taken together, our studies identify SUMO2 as a novel, selective vulnerability in SySa. Since SUMO2 inhibitors are currently in Phase 1/2 clinical trials for other cancers, our findings present a novel avenue for targeted treatment of synovial sarcoma.

**SIGNIFICANCE:** Our study identifies SUMO2 as a selective dependency in synovial sarcoma. We demonstrate that the SUMO2/3 inhibitor TAK-981 impairs sarcomagenesis and reverses the SS18-SSX fusion-driven oncotranscriptome. Our study indicates that SUMO2 inhibition may be an attractive therapeutic option in synovial sarcoma.

## INTRODUCTION

Synovial sarcoma (SySa) belongs to a subcategory of sarcomas called soft-tissue sarcomas which accounts for 5% - 10% of all soft-tissue tumors^1^ and is more prevalent in adolescents and young adults^2^. Approximately 30% of SySa cases occur in patients under twenty years of age^3,4^. This disease is characterized by an oncogenic fusion protein SS18-SSX formed by the translocation of (X;18)(p11.2;q11.2)^5,6^, which leads to the fusion of the SS18 gene to one of three SSX genes (SSX1, SSX2 or rarely to SSX4) on chromosome X. Although the SS18-SSX fusion has been characterized for more than three decades, therapies that target this fusion, or the oncogenic program driven by these fusion proteins remain to be identified.

The SS18-SSX fusion protein interacts with the SWI/SNF (BAF) complex, a large, chromatin modifying complex dysregulated in many human cancers. This interaction displaces the full-length SS18 as well as the SMARCB1/BAF47 protein from the BAF complex, altering its normal composition and function^7^. The modified BAF complex then colocalizes with the Polycomb Repressive Complex 2 (PRC2)^8^, leading to dysregulated transcriptional changes that are important for the oncogenesis of synovial sarcoma. This aberrant interplay between the BAF and PRC complexes results in the upregulation of several oncogenic pathways, including the Wnt/β-catenin^9,10^, FGFR^11^, and NOTCH^12^ pathways, while downregulating tumor suppressors such as EGR1^13,14^ and copy number variations of CDKN2A^15^ to name a few.

The SS18 in the fusion is part of the canonical BAF complex and is associated with transcriptional activation. However, the SSX portion of the fusion protein is known to be repressive in function and binds regions rich in H2AK119ub1^16^, deposited by the non-canonical PRC1.1 complex. Although the SSX portion does not contain a direct ubiquitin binding site, recent findings indicate that it specifically binds to H2AK119ub-decorated sites via the ‘H3-H2AK119ub’ basic groove^17^. This abnormal interaction leads to the unraveling of the nucleosome, redirecting the BAF complex to regions of chromatin occupied by the polycomb complex, which is one of the key mechanisms responsible for the epigenetic rewiring that drives synovial sarcoma pathogenesis.

Given the lack of targeted treatments in synovial sarcoma, a systematic approach to identify clinically tractable dependencies may yield valuable new candidates for therapy. Functional genomic approaches such as RNAi and CRISPR–Cas9 screens are powerful tools for forward genetics and have been effectively employed for the unbiased discovery of factors important for the viability of cancer cells^18–21^. These large-scale screens can be used to identify vulnerabilities that are selectively essential for certain mutational subtypes (such as BRAF or KRAS mutated cancers)^22^, or to nominate candidate targets selectively required for cancer types of interest^23^. In this study, through an analysis of the DepMap RNAi and CRISPR datasets, we identified genes that are selectively essential in SySa cell lines compared to other cancer cell lines. Custom pooled screens of the top SySa selective vulnerabilities revealed the small ubiquitin-like modifier 2 (SUMO2) as one of the most significant dependencies both *in vitro* as well as *in vivo*. Importantly, small molecule inhibition of SUMO2 using TAK-981, a mechanism-based inhibitor of the SUMO2-activating enzyme (SAE) specifically led to a diminution of the fusion protein expression, chromatin occupancy, concomitant reversal of the genetic and epigenetic “lesions” characteristic of the SySa fusion proteins and strongly impaired SySa pathogenesis *in vitro* and *in vivo*. Taken together, our results reveal SUMO2 inhibition as an attractive therapeutic strategy in synovial sarcoma.

## RESULTS

### Analysis of functional genomic screens identifies novel and known genetic vulnerabilities in synovial sarcoma

To identify potential genetic dependencies selective to synovial sarcoma, we analyzed gene dependency data from DepMap RNAi as well as CRISPR-Cas9 screen datasets and selected genes that have a higher essentiality in SySa compared to other cell lines (Fig. 1A-C). The list of SySa-selective dependencies identified through this analysis included SS18 and SSX genes that constitute the pathogenic fusions in SySa, as well as targets that have been proposed and validated by other groups including PCGF3 and BRD9^24^ (Fig. 1A-C). Our analysis also revealed several candidate SySa-selective genes that have not hitherto been studied in the context of SySa pathogenesis (Fig. 1A-C and Table S1). From the synovial sarcoma cell lines represented in the DepMap database, we selected top 200 genes from each of the datasets based on their DEMETER2 (RNAi) and Chronos (CRISPR) scores. From these lists, 351 unique genes were selected (Table S1). We then conducted pathway analysis using Enrichr^25^ to identify potential enrichment for biological pathways in the SySa-selective dataset. This analysis revealed that there was a striking enrichment for the SUMO conjugation and SUMO transfer Reactome pathway (adjusted p values of 0.03 and 0.009 respectively) and multiple members of the sumoylation machinery appeared as hits in the SySa-selective dependencies dataset including UBA2, SAE1, UBE2I, SUMO2 and, PIAS1 (Table S2 and Fig. 1D).

**Figure 1:**
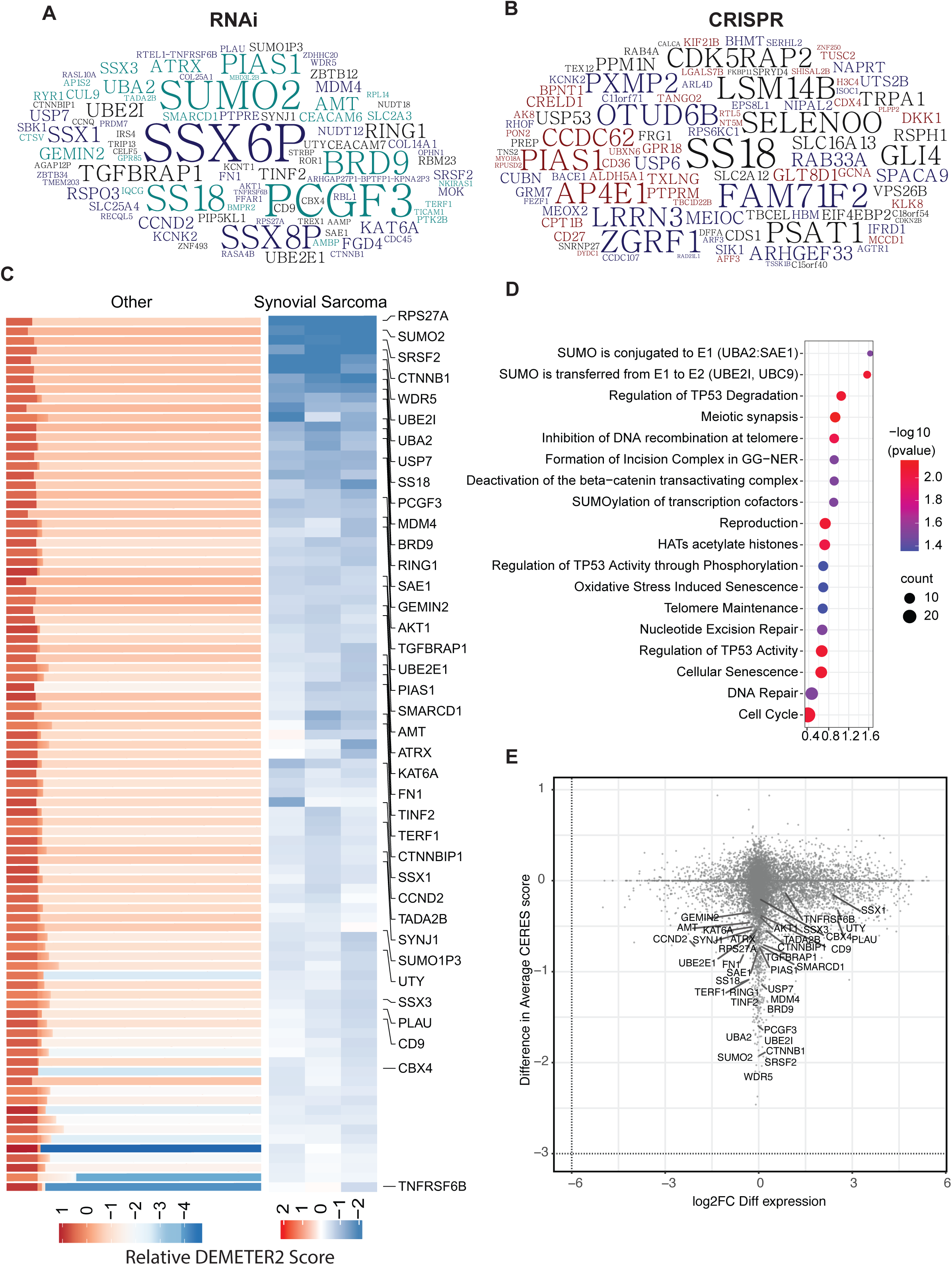
Synovial sarcoma dependencies identified through DepMap screens: **A-B)** Top 200 genes identified as selective dependencies using T-statistic scores of **(A)** RNAi data or **(B)** CRISPR screening data are shown in the word cloud. Font size is proportional to the negative log 10 adjusted p value with a larger font indicating a higher dependency of the gene in synovial sarcoma cell lines compared to all other cell lines in the DepMap database. **C)** Heatmaps representing DEMETER2 scores, as a quantitative dependency metric of human synovial sarcoma cell lines to each gene in RNAi screens. Relative DEMETER2 scores for non-synovial sarcoma cell lines (left) compared to synovial sarcoma cell lines (right) are depicted for the top differentially essential genes. **D)** Bubble plot displays the top significantly enriched pathways in the REACTOME database with adjusted p values < 0.05. **E)** Scatter plot showing the relationship between gene dependency (measured by the difference in average DEMETER2 scores) and differential transcript expression (log2 fold change FC differential expression) for all genes, with key synovial sarcoma selective essential genes labeled.

Other biological pathways enriched in this SySa-selective dependency data included genes involved in meiotic synapse formation, deactivation of the beta−catenin transactivating complex, histone acetylation, and regulation of p53 activity. Analysis of these SySa-selective dependencies using the STRING database showed enrichment in protein complexes involved in chromosome organization, WNT signaling, BAF complex, and PRC1 activity (Fig. S1) which are known dependencies in synovial sarcoma^7,17,24,26^. Novel biological pathways and protein complexes identified included the SUMO2-UBE2I complex, the SAGA, and the synaptojanin complex (Fig. S1). Next, we wanted to evaluate whether genes selectively essential for SySa were differentially expressed at the transcriptional level in SySa cell lines compared to other cancer cell lines. Thus, we calculated the fold change for each of these genes between SySa and non-SySa cancer cell lines in the Cancer Cell Line Encyclopedia (CCLE) and plotted it against the relative dependency values (DEMETER2) (Fig. 1E). In this analysis, we observed that while genes such as SSX1 and SSX3 were indeed much more highly expressed in SySa compared to non-SySa cell lines, genes such as BRD9, PCGF3, and SUMO2 had no noticeable difference in expression between these cell lines (Fig. 1E). This analysis indicates that while the relatively higher dependency of SySa cell lines on the SSX genes may result from their higher expression in cell lines from this lineage compared to others, the dependence on genes such BRD9, PCGF3, and SUMO2 may instead be explained by a relatively higher activity of these proteins in SySa compared to other cancers.

### *In vivo* and *in vitro* CRISPR screens nominate new candidate targets in synovial sarcoma

Building on our previous analysis, we sought to test these SySa-selective dependencies more comprehensively and investigate their essentiality in an *in vitro* as well as *in vivo* setting. To do so, we set up pooled CRISPR/Cas9 screens for the SySa-selective genes. First, we assessed the activity of Cas9 in HS-SY-II cells expressing Cas9 to ensure high editing efficiency (indel percentage identified as ∼ 92% and a knockout score of 90 using ICE^27^). With these optimized conditions, we then performed parallel in *vivo* and *in vitro* CRISPR screens (schematic Fig. 2A). HS-SY-II-Cas9 cells expressing Cas9 were transduced with the screening library in duplicate at a MOI of ∼0.3. We then subcutaneously injected 2 million cells (∼500X coverage) into the flanks of nude mice. In parallel, for the *in vitro* screen, we cultured the cells from each replicate for ∼10 doubling times. There was strong replicate reproducibility for both the i*n vitro* and *in vivo* results (Fig. S2). sgRNA abundance and distribution were quantified using MAGeCK Robust Rank Aggregation algorithm^28^. *In vitro* and *in vivo* hits were generally well correlated (Fig. S2), with the identification of a number of overlapping hits including KAT2A, C8orf82, SUMO2, FRG2, BICDL1, and LGALS7B (Fig. 2B-D and Table S3). We then turned our attention to targets that were previously not described as dependencies of synovial sarcoma and ranked highly in both the *in vivo* as well as *in vitro* screens (Fig. 2D). To further prioritize these hits, we also overlapped them with genes that are regulated by the SS18-SSX fusion oncoprotein (SS18-SSX fusion targets) as shown by Jerby-Arnon et.al^29^ (Fig. 2E) (Table S3). Of the genes that are strongly depleted in our *in vitro* and *in vivo* screens and are activated by SS18-SSX fusions in SySa cells, we were particularly interested in SUMO2. SUMO2 was one of the most essential genes in the *in vitro* (RRA score 5.29E-06), as well as the *in vivo* screen (RRA score 7.95E-05). Interestingly, SUMO2 has been shown to be transcriptionally activated by SS18-SSX fusions in two independent SySa cell lines in prior studies^29^. Pathway enrichment analysis showed that the top hits were enriched for proteins involved in the SUMO complex in both *in vitro* as well as *in vivo* screens (Fig. 2F-G). Individual sgRNAs for SUMOylation pathway genes showed a dramatic drop in read counts (Fig. 2H-J) further validating SUMO2 as a top candidate hit in our screens. A small molecule inhibitor - TAK-981, that selectively inhibits SUMO2 is currently in phase 1/2 clinical trial for Non-Hodgkin lymphoma (NCT04074330) and phase 1b/2 for refractory multiple myeloma (NCT047760180). We therefore earmarked SUMO2 as a novel candidate and a therapeutic target for further evaluation.

**Figure 2:**
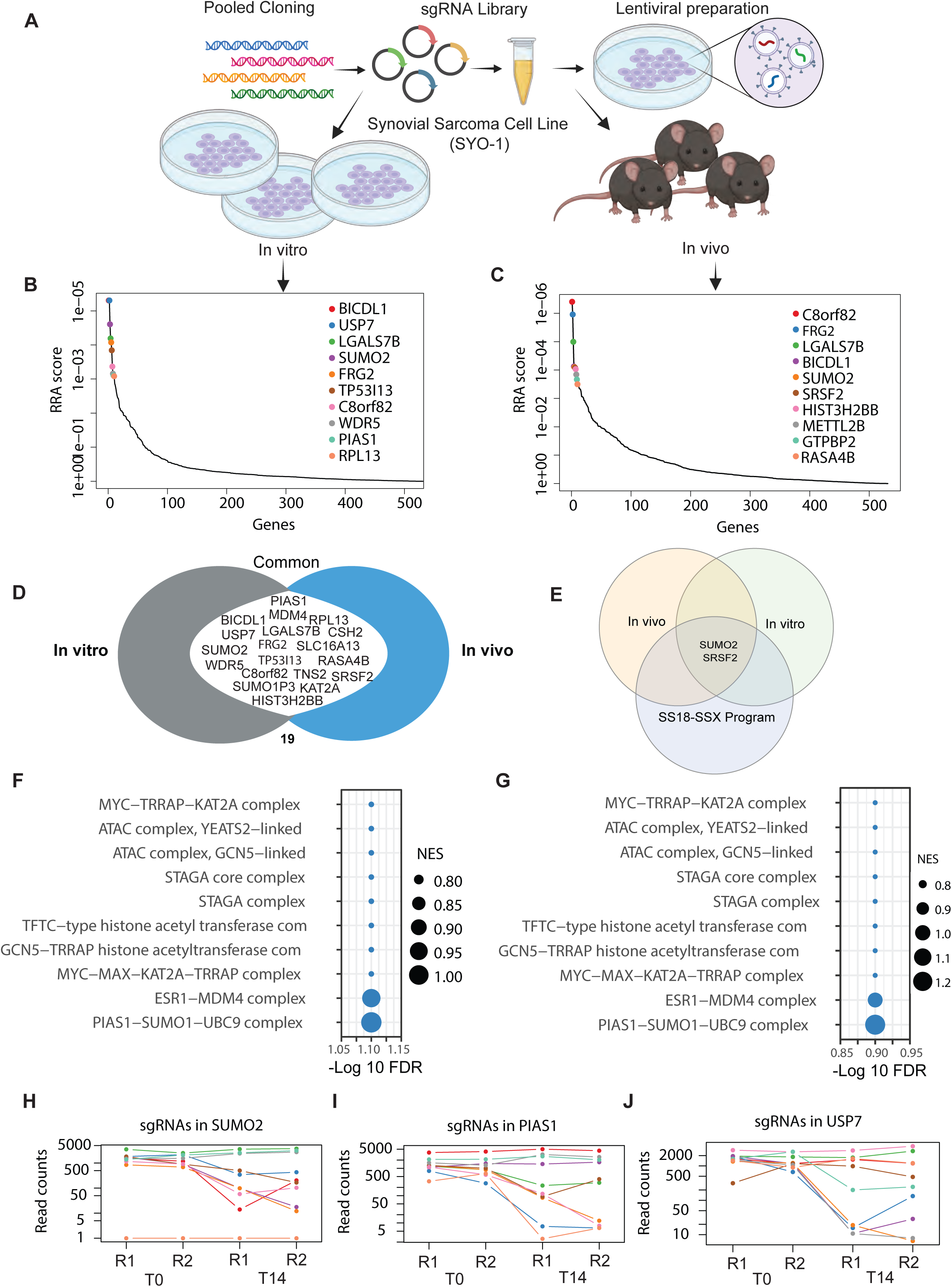
*In vivo* and *in vitro* screening reveal top synovial sarcoma-selective dependencies: **A)** Schematic representation of *in vivo* and *in vitro* pooled CRISPR screens in HS-SY-II cell line. **B-C)** Analysis of pooled *in vitro* (left) and *in vivo* (right) CRISPR/Cas9 screens using the MAGeCK RRA algorithm. Plot shows the relationship between genes (X-axis) and their statistical significance (RRA score) (Y-axis). The top 10 significant genes are labeled. RRA is robust rank algorithm as assessed using MAGeCK. **D)** A Venn diagram illustrating genes commonly essential in both the *in vitro* and *in vivo* pooled CRISPR/Cas9 screens. Common genes in the union are labeled. **E)** A Venn diagram illustrating genes common to the *in vitro* and *in vivo* screens as well as in the core synovial sarcoma oncogenic program. Common genes in the union are labeled. **F-G)** Pathway enrichment of hits in the in vitro (**F**) and in vivo screens (**G**) screens. -log10(FDR) of false discovery rate is shown on the X-axis. The size of the bubbles indicates normalized enrichment scores of each pathway. **H-J)** Normalized read counts of multiple individual sgRNAs for SUMO2 **(H)**, PIAS1 **(I)** and USP7 **(J)** showing the difference between T0 and T14 time points.

### TAK-981, a small molecule SUMO2 inhibitor impairs the growth of synovial sarcoma cells

To systematically test the effect of SUMO2 inhibition on synovial sarcoma cell lines, we first evaluated the effect of TAK-981 on proliferation in four different human synovial sarcoma cell lines (SYO1, HS-SY-II, 1273/99, Aska-SS) as well as the epithelial squamous cell lung cancer cell line (SK-MES-I) and human embryonic kidney 293T cells (HEK-293T). TAK-981 treatment diminished sumoylation (Fig. S3) and significantly reduced the proliferation of these cell lines in a concentration-dependent manner, showing half maximal effective concentration (EC_50_) in the nanomolar range in a CellTiter-Glo assay, with the HS-SY-II cell line exhibiting the strongest inhibition (Fig. 3A). Generally, SySa cells lines showed a substantially higher sensitivity to TAK-981 than SK-MES-I or HEK293-T cells (Fig. 3A). To determine the effect of TAK-981 on apoptosis, we performed Annexin V staining - on TAK-981 treated and untreated cells. The proportion of early apoptotic cells significantly increased in HS-SY-II and SYO1 cells after 24 hours of TAK-981 treatment compared to DMSO-treated cells (Fig. 3B and 3C). Additionally, cell cycle analysis using propidium iodide indicated an S-phase arrest (Fig. 3D). We then performed cell viability assays on 2D and 3D cultures for SYO1 and Aska-SS cell lines to determine whether these culture conditions resist TAK-981 treatment. In these studies, too, TAK-981 treatment led to a progressive and marked decrease in viability as measured by CellTiter-Glo in both 2D as well as 3D cultures conducted over 2, 3, and 4 days (Fig. 3E and 3F). Colony-forming assays for the SYO1, HS-SY-II and 12273/99 cell lines using a TAK-981 titration series also demonstrated a dramatic and dose-dependent reduction in colony formation (Fig. 3G).

**Figure 3:**
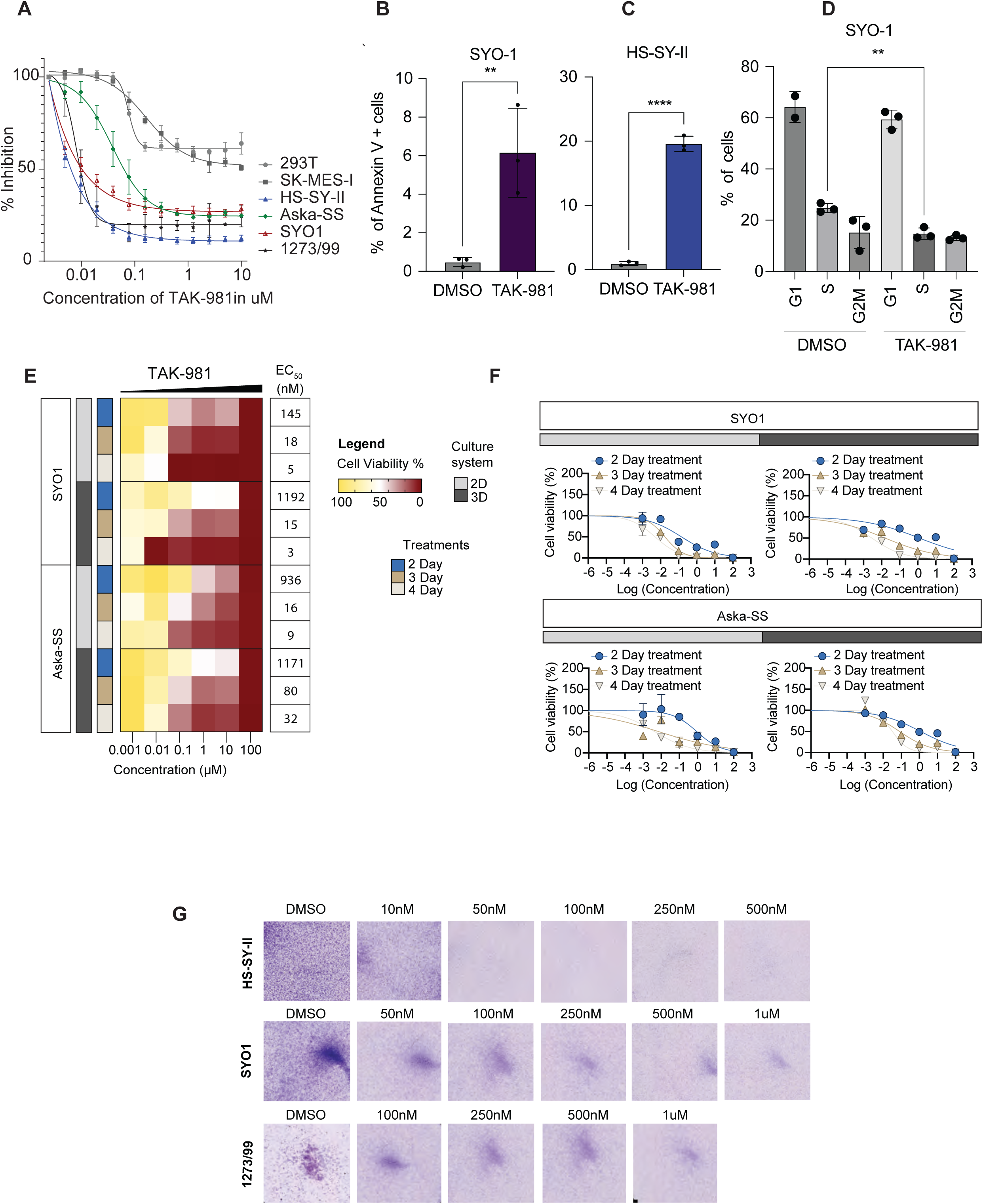
Effect of TAK-981 on synovial sarcoma cell lines: **A)** Viability of various SySa cell lines measured using Cell-Titer-Glo after 48 hrs of treatment with varying concentrations of TAK-981 is shown. X-axis shows concentration of TAK-981 and Y-axis shows percent inhibition compared to vehicle-treated counterparts. N=4. **B-C)** Percent Annexin V positive SYO1 and HS-SY-II cells in the DMSO compared to TAK-981 treated arms are plotted on the Y-axis. N=3, 48 hrs of treatment with p-value legend ^∗∗^, *P* < 0.01; ^∗∗∗^, *P* < 0.001; \*\*\*\**P* < 0.0001. **D)** Percent propidium iodide (PI) positive SYO1 cells in DMSO-treated compared to TAK-981-treated arm are shown (Y-axis). N=3, 48 hrs post treatment with p-value legend ^∗∗^, *P* < 0.01. **E)** Heatmaps denoting the viability as a percentage of vehicle-treated SYO1or Aska-SS cells treated with TAK-981 in varying concentrations (indicated on the X-axis) in 2D and 3D cell culture formats for days 2, 3 or 4 are shown. The right column indicates the EC50s in each condition. Legends for the cell culture system used or the treatment day are shown in the center. **F)** Percent viability (relative to DMSO-treated) SYO1 (top) or Aska-SS cells (bottom) 2,3 or 4 days after treatment in 2D or 3D growth formats is plotted (Y-axis). **G)** Pictures showing crystal violet stained colonies in HS-SY-II, SYO1 or 1273/99 cell lines 48hrs after treatment with DMSO or varying concentrations of TAK-981.

### TAK-981 treatment impairs transcription of key oncogenic pathways in synovial sarcoma cell lines

To comprehensively interrogate the transcriptomic changes occurring in synovial sarcoma cells upon TAK-981 treatment, we treated HS-SY-II (harboring the SS18-SSX1 fusion) and SYO1 cells (harboring the SS18-SSX2 fusion) with DMSO or TAK-981 and performed bulk RNA sequencing. Common to both HS-SY-II and SYO1, a total of 1100 differentially expressed genes (DEGs) were detected using the threshold of |fold change| >2 and adjusted *p*-value < 0.01, of which 908 and 192 genes were upregulated or downregulated, respectively (Table S4). Of these, key cancer-associated genes shown to be upregulated by the SySa fusion^29^ were downregulated by TAK-981 treatment, including *CDX2*, *HOXA10*, *SUZ12*, *TYMS*, *AURKB*, (Fig. 4A) and *HOXC10* and *SMC2* (Fig. 4B). Concomitantly, genes upregulated by the SySa fusions were downregulated by TAK-981 treatment including *KLF4*, *GADD45B*, *CXCR4* and *GDF15* (Fig. 4A-B). The commonly downregulated genes were highly enriched for cell cycle (adjusted P value 1.023e-39), cell cycle checkpoint (adjusted P value 2.000e-22) and S phase genes (adjusted P value 2.304e-17), and DNA replication-associated genes (adjusted P value 4.018e-13) in the Reactome database, consistent with cell cycle arrest following SUMO2 inhibition. Importantly, it has been shown that the high expression of cell cycle genes is a key feature of a subset of undifferentiated cells in synovial sarcoma patient samples and that these genes are regulated by SS18-SSX fusions. In our studies, these genes showed a significant downregulation upon TAK-981 treatment (Fig. 4C). Of genes that were commonly upregulated by TAK-981 treatment in the two cell lines, there was a significant enrichment of genes involved in collagen formation and extracellular matrix formation (adjusted P value 3.435e-8). Notably, TAK-981 treatment also led to the significant downregulation of several genes associated with resistance to doxorubicin, which is used in the treatment of synovial sarcoma^30^ Fig. 4C-D) as assessed using gene set enrichment analysis (GSEA).

**Figure 4:**
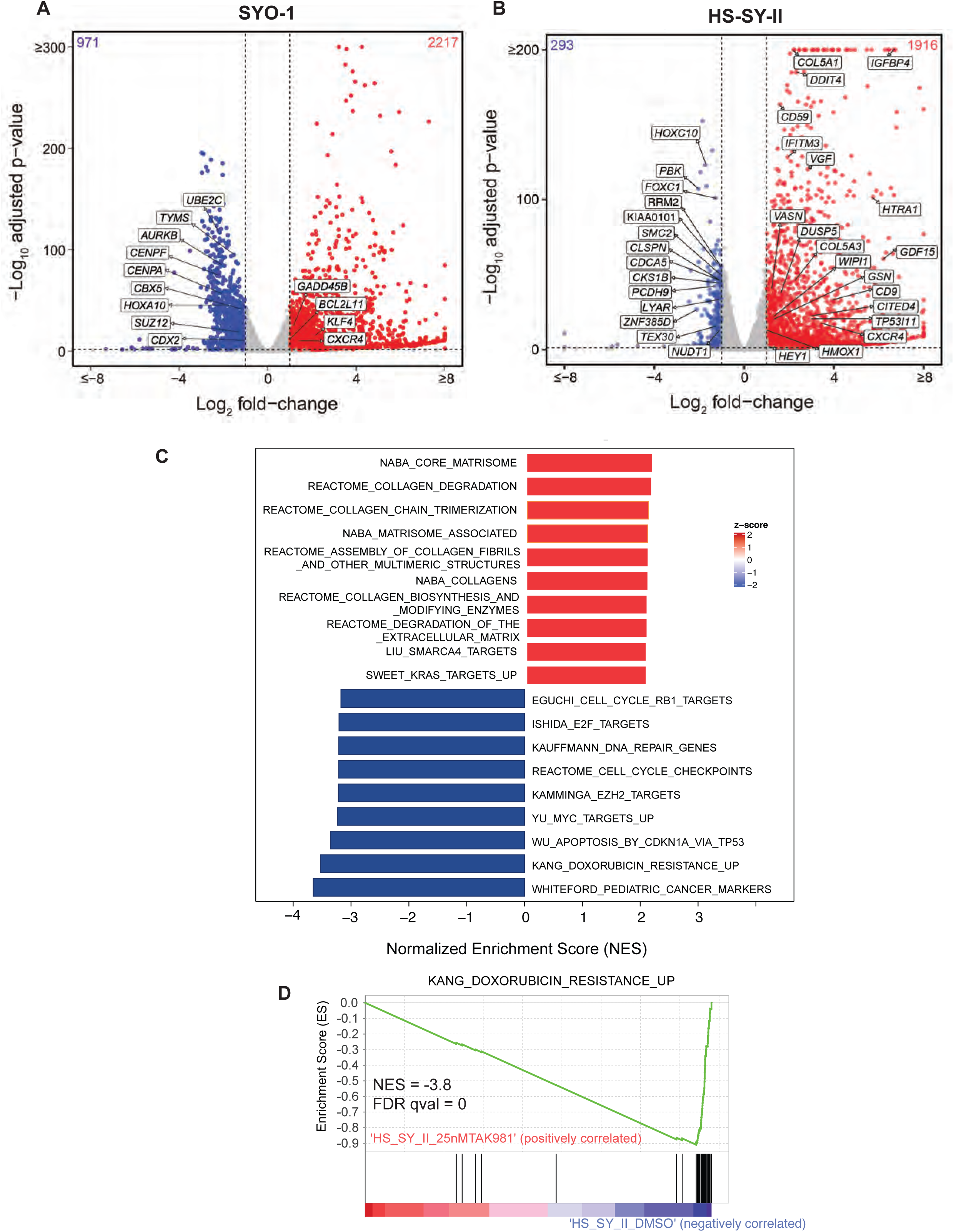
Broad transcriptomic changes in TAK-981 treated synovial sarcoma cell lines. **A-B)** Volcano plot illustrating differential gene expression in SYO1 **(A)** or HS-SY-II cells **(B)** treated with DMSO compared to TAK-981. Each dot represents an individual gene. The X-axis represents log-2 fold change (DMSO Vs TAK-981 treated cells) and the Y-axis represents -log10 BH adjusted p-value. Red dots represent genes significantly upregulated in the TAK-981 treated compared to the DMSO treated arm with adjusted p-value <0.05 and fold change >2. Blue dots represent genes that are significantly downregulated with p-value <0.05 and fold change <0.5. Grey dots represent genes that are not significantly differentially expressed. **C)** The bar chart represents top 20 significantly enriched gene ontology (GO) terms of the C2: canonical pathways gene set. The horizontal axis represents pathways with positive (red) and negative (blue) normalized enrichment scores (NES). **D)** GSEA analysis of the C2 Curated Datasets in MiSigDB for genes upregulated during doxorubicin resistance is shown for transcriptomic data of HS-SY-II cells treated with DMSO compared to TAK-981. NES = Normalized Enrichment Score.

### TAK-981 treatment specifically reverses the transcriptional signatures driven by SySa fusion proteins

Next, we investigated whether TAK-981 treatment specifically alters the expression of synovial sarcoma fusion target genes, as defined by Jerby-Arnon et al^29^. In their study, the authors defined the SS18-SSX program by knocking down the SS18-SSX fusion and conducting a ChIP-seq analysis for the fusion. This allowed them to identify genes that were bound by the SS18-SSX fusion protein and whose expression was modulated by the knockdown of the fusion as direct targets and genes not bound by the fusion but modulated by its knockdown as indirect targets. We utilized this list of genes for a custom gene set enrichment analysis in the TAK-981 treated RNA-seq dataset.

In these analyses, we observed that in both SYO1, and HS-SY-II cell lines, TAK-981 treatment led to a dramatic reversal of the SS18-SSX-driven transcriptomic program. Specifically, genes activated by the chimeric SS18-SSX fusion protein showed a significant reduction in expression upon TAK-981 treatment as assessed using GSEA (Fig. 5A-B), and this included the HOX genes HOXC6, HOXA10 as well as SRSF1 and TYMS (Fig. 5C). Concomitantly, genes repressed by SS18-SSX in synovial sarcoma were reactivated (Fig. 5C-D), including KLF4, TBX3 and CXCR4 genes (Fig. 5E). Of note, the fact that this was evident both in the SYO1 cell line expressing the SS18-SSX2 fusion protein as well as in the HS-SY-II cell line expressing the SS18-SSX1 fusion strongly indicate that SUMO2 is critical for the transcriptional activity of both types of distinct SS18-SSX fusion oncoproteins.

**Figure 5:**
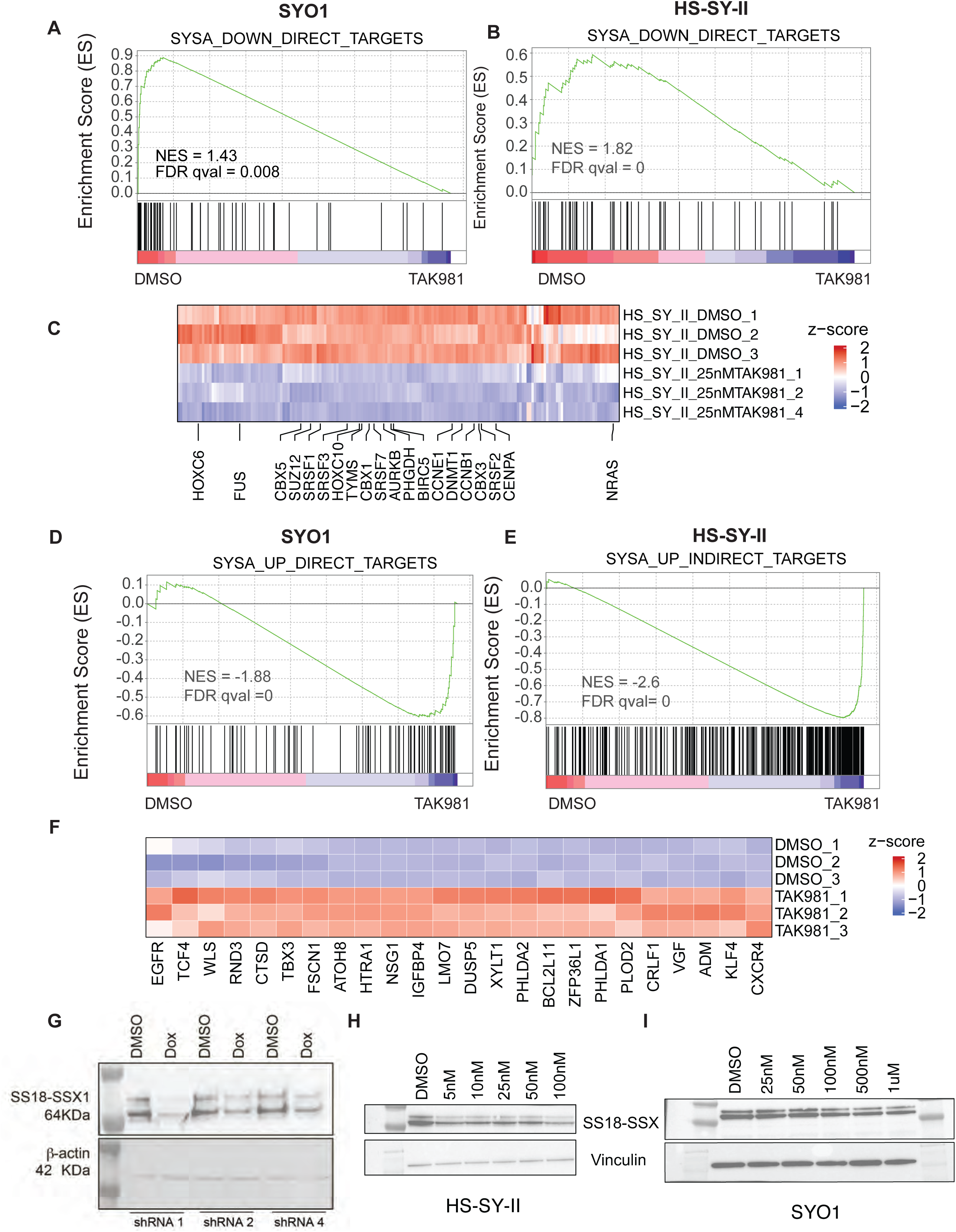
Treatment with TAK-981 leads to downregulation of oncogenic program in synovial sarcoma cell lines. **A-B)** GSEA analysis of SS18-SSX fusion-activated genes in SYO1 (**A**) or HS-SY-II (**B**) cells treated with DMSO compared to TAK-981 is shown. Enrichment plots depict genes that are direct targets of the SS18-SSX fusion, which are downregulated upon SS18-SSX fusion knockdown (thus, SS18-SSX-activated genes). Black vertical lines at the bottom indicate positions of individual genes in the set, with the green line representing the cumulative enrichment score (y-axis). A positive normalized enrichment score (NES) indicates enrichment in the upregulated genes in SYO1 cells **(A)** and HS-SY-II cells **(B).** FDR q values are indicated. **C)** Heatmap displaying SS18-SSX fusion-activated genes that are reduced in TAK-981 treated compared to DMSO treated HS-SY-II cell line are shown. Select genes implicated in SySa pathogenesis are labeled. **D-E)** GSEA analysis SS18-SSX fusion-repressed genes in SYO1 (**D**) or HS-SY-II (**E**) cells treated with DMSO compared to TAK-981 is shown. Enrichment plots show genes that are indirectly repressed by the SS18-SSX fusion and are thus upregulated upon SS18-SSX fusion knockdown. A negative NES indicates higher expression enrichment of these genes in the TAK-981 compared to DMSO arms in SYO1 **(D)** as well as HS-SY-II cells **(E)**. **F)** Heatmap displaying SS18-SSX fusion-repressed genes that are increased in TAK-981 treated compared to DMSO treated HS-SY-II cell line are shown. Select genes implicated in SySa pathogenesis are labeled. **G)** Immunoblot analysis of whole-cell lysates from HS-SY-II cells stably expressing SUMO2 knockdown in a doxycycline-inducible system (shRNA1, 2, and 4), probed for the SS18-SSX1 fusion protein with Vinculin as a loading control is shown. **H-I)** Immunoblot analysis of whole-cell lysates from HS-SY-II **(H)** or SYO1 cells **(I)** treated with varying denoted concentrations of TAK-981 and probed for the SS18-SSX1 fusion protein with are shown. Vinculin is shown as a loading control.

Our observation that SUMO2 inhibition reverses the oncogenic program driven by two distinct SS18-SSX fusion oncoproteins indicates that SUMO2 is a critical node in regulating the oncogenic activity of these chimeric oncoproteins. Given the specific reversal of the SySa fusion-driven program, we sought to test the intriguing hypothesis that SUMO2 regulates the SS18-SSX fusion protein itself. For this, we cloned shRNAs targeting SUMO2 into a tetracycline-inducible plasmid and expressed the shRNAs in HS-SYII cells. qPCR results validated the knockdown of SUMO2 transcript expression following doxycycline induction of the shRNAs (Fig.S4). Strikingly, SUMO2 knockdown with 3 independent shRNAs showed a dramatic reduction in SS18-SSX1 protein levels in HS-SYII cells (Fig. 5G). These results could be replicated using a pharmacologic approach - TAK-981 treatment led to a reduction in the levels of SS18-SSX1 protein in the HS-SY-II cell line (Fig. 5H) and SS18-SSX1 fusion in the SYO1 cell lines (Fig. 5I) as well as in the 1273/99 cell line (Fig. S5). These results provide a striking demonstration that SUMO2 inhibition modulates the levels of oncogenic fusion proteins that drive sarcomagenesis in SySa.

### SUMO2 inhibition diminishes SS18-SSX chromatin occupancy and reverses fusion-driven aberrant epigenomic changes in SySa cells

Next, we sought to assess whether TAK-981 treatment affects the chromatin occupancy of the SS18-SSX fusion protein. For this, we performed Cleavage Under Targets & Release Using Nuclease (CUT&RUN) using the SS18-SSX-fusion specific antibody (see Methods). These studies demonstrated a substantial decrease in SS18-SSX2 fusion genomic occupancy as assessed using spike-in normalized CUT&RUN analysis in the TAK-981-treated compared to vehicle-treated arms (Fig. 6A). Specifically, TAK-981 treatment of SYO1 cells showed a 1.87-fold reduction in genome-wide chromatin binding signal of the SS18-SSX fusion compared to the DMSO treated cells, as computed from fraction of reads in peaks (FRiP) measured using consolidated peaks in DMSO replicates. A meta-analysis of the fusion-binding signal at synovial sarcoma target genes^29^ revealed a reduction in the fusion binding with the maximum signal centered around the transcription start site (Fig. 6B). Since increased H2AK119ub deposition has been linked to the pathogenic activity of the SS18-SSX fusions, we then sought to assess H2AK119 ubiquitination in TAK-981 treated cells. Interestingly, our studies showed a marked reduction in H2AK119ub in SYO1, HS-SY-II, 1273/99 and Aska cell lines treated with TAK-981 as assessed using immunoblotting (Fig. 6C). Furthermore, chromatin immunoprecipitation (ChIP)-sequencing of H2AK119ub showed that similar to the loss of SSX-SS18 expression, there was a substantial reduction in H2AK119ub in TAK-981 compared to DMSO-treated cells genome-wide (Fig. 6D). Specifically, there was a 1.53-fold reduction of genome-wide H2AK119ub levels in TAK-981 versus DMSO treated SYO1 cells, computed as fraction of reads in peaks (FRiP) measured using consolidated peaks in DMSO replicates. Genes including the SS18-SSX-activated targets HOXA10 and SOX8 lost SS18-SSX occupancy and showed reduced expression upon TAK-981 treatment (Fig.6F and Fig. S6). Concomitantly, SS18-SSX-repressed targets such as GADD45B showed diminished SS18-SSX fusion occupancy, and reduced H2AK119ub, and showed increased expression (derepression) following TAK-981 treatment (Fig.6F). These results further reinforce the notion that TAK-981 treatment reverses the transcriptional activity of the pathogenic SS18-SSX fusion.

**Figure 6:**
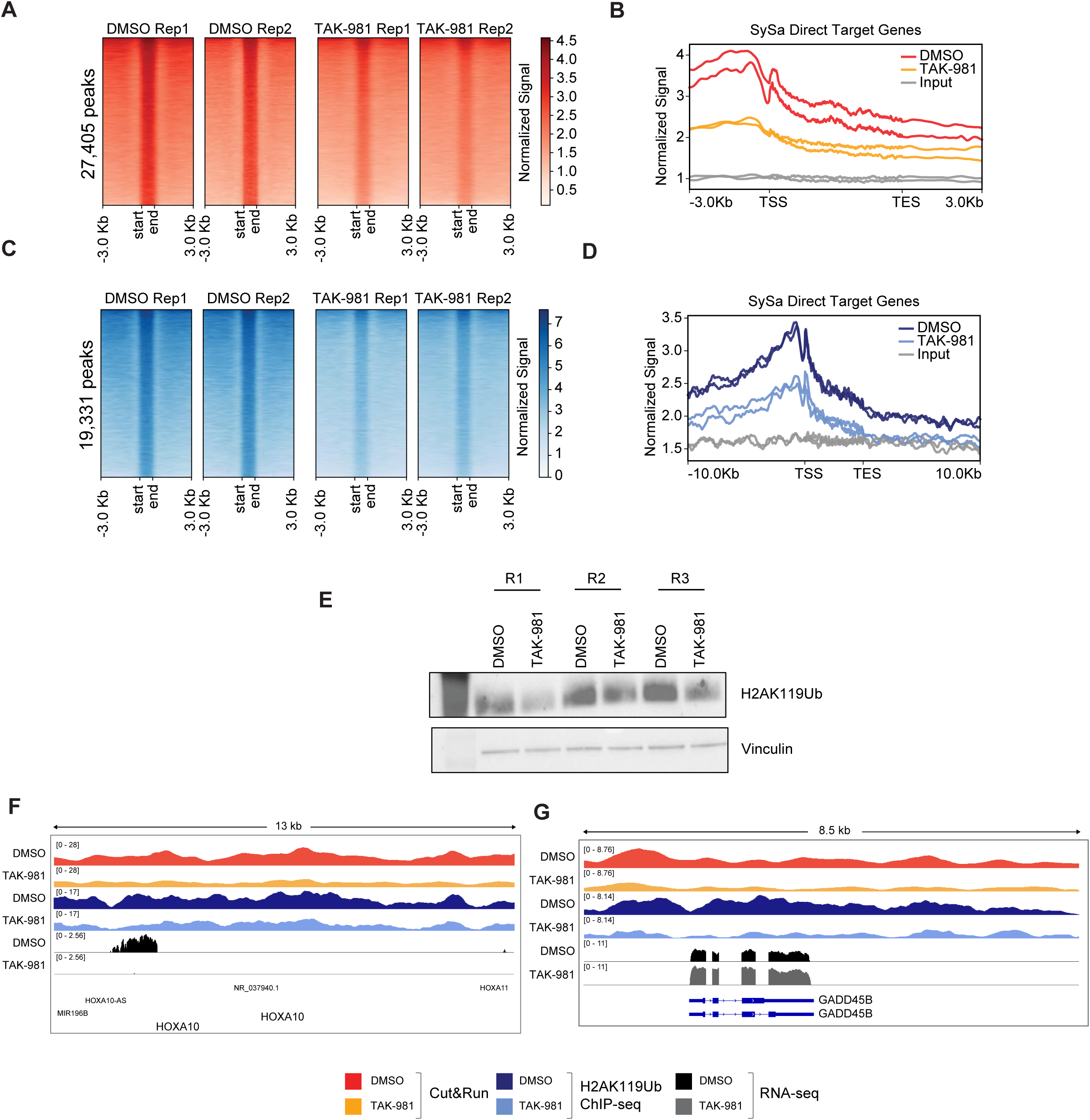
Treatment with TAK-981 causes the fusion oncoprotein SS18-SSX1 eviction from chromatin. **A)** Cut&Run density heatmaps of SS18-SSX occupancy in SYO1 cells treated with DMSO (left) or TAK-981 (right) across 27,405 peaks are shown. N=2 **B)** Meta-analysis plot showing normalized SS18-SSX binding signal (Y-axis) at gene bodies from the transcription start site (TSS) to the transcription end site (TES) centered around the TSS +/− 3Kb for SySa direct target genes is shown. **C)** CHIP-seq density heat maps of H2AK119Ub occupancy in SYO1 cells treated with DMSO or TAK-981 for 72 hrs across 19,331 peaks is shown. **D)** Meta-analysis plot showing normalized H2AK119 signal at the genic loci of SySa direct target genes +/− 3Kb is shown. **E)** Immunoblot of H2AK119ub on SYO1 cells treated with TAK-981 compared to DMSO control are depicted. Vinculin is shown as a loading control. N=3. **F)** Integrated genome viewer (IGV) tracks for the SS18-SSX fusion and H2AK119ub in DMSO or TAK-981 treated SYO1 cells along with corresponding RNAseq tracks are shown for genes HOX10A (**F**) and GADD45B (**G**).

### TAK-981 impairs sarcomagenesis of SySa *in vivo*

To determine the antitumor activity of TAK-981 *in vivo*, we injected SYO1 (harboring the SS18-SSX2 fusion) or Aska-SS cells (harboring the SS18-SSX1 fusion) into the flanks of nude mice. When tumors became palpable, mice were treated with 25mg/kg of TAK-981 or vehicle. A dosing schedule of 3 intraperitoneal injections a week for 5 weeks was maintained (Fig. 7A). Consistent with the *in vitro* assays, TAK-981-treated mice showed a remarkable reduction of tumor growth when compared to vehicle-treated mice. Tumor volumes in Aksa-SS injected mice were significantly reduced in the TAK-981-treated arm as were tumor weights (Fig. 7B-D). IHC analysis of the tumors stained with hematoxylin and eosin showed a marked reduction in the number of cells per unit area within TAK-981 treated tumors when compared to the vehicle-treated tumors (Fig. 7E-I), both in the periphery and center of the tumor (Fig. 7E-F). Ki67 staining revealed a ∼60% decrease in Ki67 positivity in comparison with the vehicle-treated tumors indicating decreased proliferation (Fig. 7G-I). We observed that TAK-981 was well tolerated, and the mice maintained their body weight and showed no visible signs of toxicity through the dosing period (Fig. S7). Similar results were obtained for SYO1 injected mice, where TAK-981 treatment led to a significant decrease in tumor size (Fig. 7J-L), and a concomitant decrease in cellularity (Fig. 7M&N) and Ki67 positive cells (Fig.7O-Q). The data demonstrates that TAK-981 efficiently inhibits tumor growth in SS18-SSX1 fusion containing ASKA-SS as well as SYO1 cell lines. Taken together, these data demonstrate that TAK-981 treatment has potent *in vivo* activity *in in vivo* models of SySa tumors.

**Figure 7:**
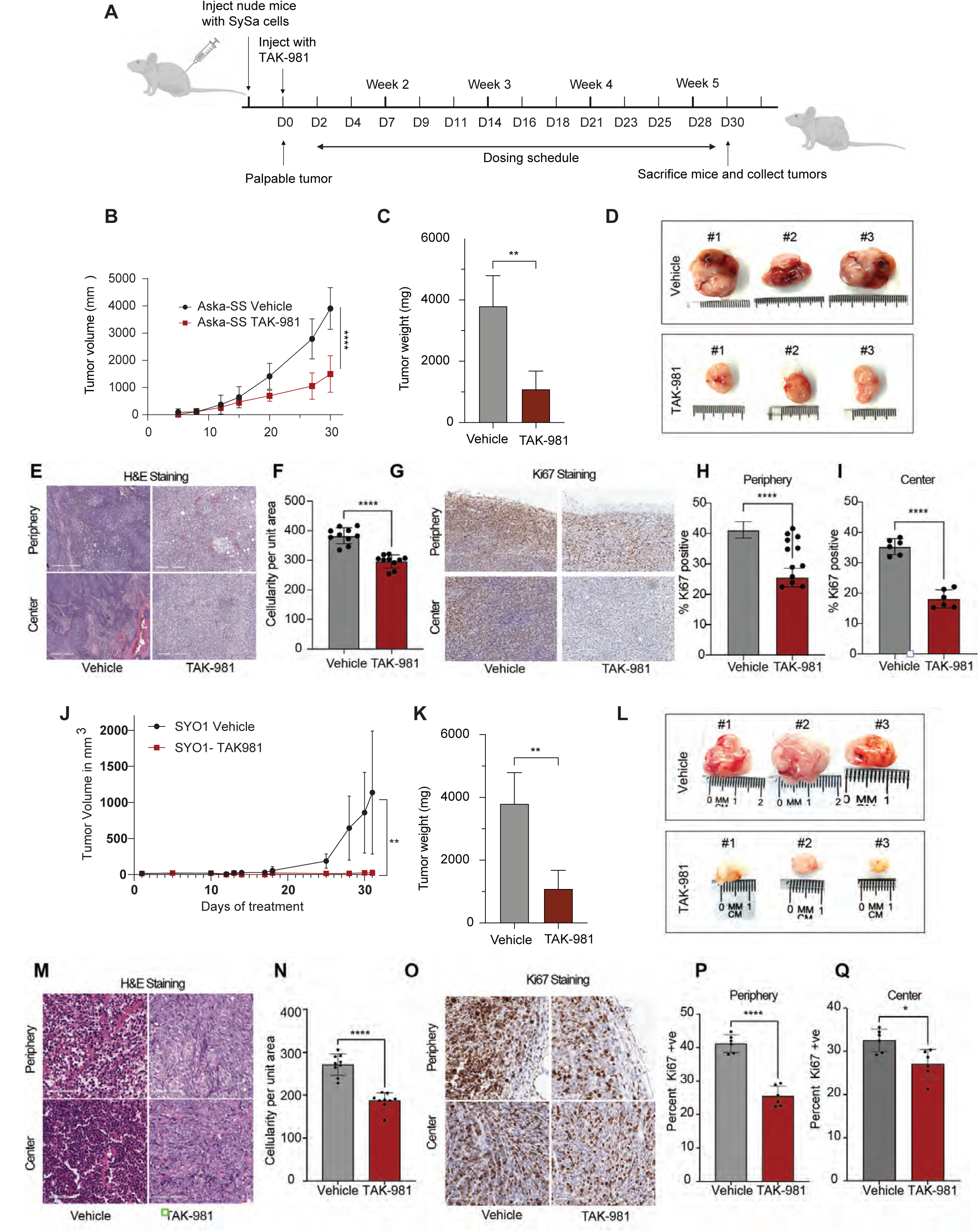
Tumor size reduction is seen in mice treated with TAK-981: **A)** Schematic showing the *in vivo* experiment with TAK-981 treatment in Aska-SS or SYO1 injected nude mice. The cartoon depicts time of cell injection, duration and frequency of treatment, and time of final tumor harvesting. **B)** Average tumor volumes of mice injected with Aska-SS are shown for the duration of the experiment. **C)** Tumor weights for DMSO or TAK-981 treated Aska-SS mice are shown. N=3 **P<0.01, student’s t-test. **D)** Representative images of extracted Aska-SS tumors **E)** Immunohistochemical staining of tumors with hematoxylin-eosin (H & E) staining reflecting tumor areas from vehicle treated and TAK-981 treated Aska-SS-injected mice. **F)** Quantification of tumor cellularity in vehicle-treated and TAK-981-treated Aska-SS injected groups. Quantitative data represented as mean ± standard deviation (SD) for n = 10 fields per sample. **G)** Representative IHC images showing Ki67 staining in the vehicle and TAK-981 treated Aska-SS tumor sections from the periphery (top) and center (bottom) of tumors. Staining indicates Ki67-positive cells - a marker of proliferation. **H-I)** Quantification of images in G. represented as mean ± standard deviation (SD) for n = 6 fields per sample. **J-Q)** Similar results are shown for SYO-1 injected tumors, **J)** shows tumor volumes over time (N=7), **K)** shows tumor weight quantification (N=5), **L)** representative tumor images, **M)** displays H&E staining in Vehicle-treated compared to TAK-981-treated SYO-1 injected mouse tumors, **N)** shows the quantification of cellularity per unit are, while **O-Q)** show pictorial and quantitative depiction of Ki67 staining in periphery or center in tumors of SYO1 injected mice. P-value legend ^∗^, *P* < 0.05; ^∗∗^, *P* < 0.01; ^∗∗∗^, *P* < 0.001; \*\*\*\**P* < 0.0001; *P* =ns, not significant in a students T-test

## DISCUSSION

Synovial sarcoma can be managed effectively with surgery and accompanying radiation therapy and/or chemotherapy in some patients - especially in children with localized disease. However, advanced stages of the disease present with a much more difficult challenge and the prognosis in such cases remains poor. Developing more precises, targeted therapies for synovial sarcoma has been hampered by the lack of a detailed understanding of the mechanisms that drive disease pathogenesis. The presence of the disease-defining SS18-SSX protein that is largely responsible for driving tumorigenesis has prompted several efforts in trying to understand the mechanistic underpinnings of this disease. Since SS18 - the larger component of the SS18-SSX fusions – is a member of the BAF (aka SWI/SNF) chromatin remodeling complex, seminal studies sought to investigate how SS18-SSX fusions perturb normal BAF complex function. A series of studies showed that the SS18-SSX fusion protein replace the normal SS18 protein in the BAF complex, leading to the disruption of normal BAF complex activity^31^ ^7^. Further studies demonstrated that this epigenetic rewiring fundamentally alters the chromatin crosstalk between the BAF complex and the polycomb regulatory complexes PRC1 and PRC2^17,32^. Specifically, studies showed that in addition to compromising normal BAF function, the SS18-SSX containing BAF complex evicts PRC2 from fusion bound sites^32^. Lastly, more recent studies have shown that the SS18-SSX fusions enhances the activity of the PRC1 (specifically the PRC1.1) complex, through stabilization of PCR1.1 core components, enhancing global H2K119ub^17^. In fact, a recent study showed elegantly, using a conditional mouse model of SySa driven by the SS18-SSX2 fusion protein, that the H2AK119ub mark is acquired gradually during tumorigenesis, ostensibly throught the stabilization of key PRC1.1 complex members, enabling further fusion protein binding^17^. Since SS18-SSX fusions bind to H2AK119ub through the SSX reader domain that is retained in the fusion protein^33^, a picture emerges where epigenetic rewiring by the SS18-SSX fusions drive a transcriptional feed-forward loop to sustain activity of the SySa oncotranscriptome^17,26,29,34^. Considering this, it is interesting to note that SUMO2 inhibition reverses this epigenetic rewiring by reducing levels of both the SS18-SSX fusion protein as well as of global and fusion-locus specific H2AK119 ubiquitination. These results indicate that SUMO2 is likely involved in key processes that sustain the transcriptional feed-forward loop characteristic of SySa tumor cells. The effectiveness of SUMO2 inhibition in synovial sarcoma models by specifically suppressing the pathogenic features of the SS18-SSX fusion oncoprotein indicate that SUMO2 is a highly selective vulnerability in synovial sarcoma as indicated by our analysis of the Dependency Maps (DepMap) data.

Recognizing the importance of SUMO2 in other malignancies, TAK-981 - a specific SUMO2 inhibitor – has been developed for clinical testing for many solid tumors as well as hematological malignancies. Proof of concept of its efficacy has been shown in AML^35^ and pancreatic cancer^37^ in preclinical studies. However, its potential benefits in synovial sarcoma (SySa) have not been explored, and merit clinical investigation based on our findings.

Of note, SUMO2 inhibition using TAK-981 was recently shown to potentiate the antitumor immune response by activating CD8+ T-cells through modulation of type I interferon signaling^38^. In this study, TAK-981 improved the survival of mice in models of colorectal cancer, enhancing the response of anti-PD1 or anti-CTLA4 antibodies. In future studies, it will be interesting to determine whether TAK-981 treatment has similar effects on augmenting antitumor immunity in synovial sarcoma in addition to the strong cell-intrinsic anti-oncogenic activity observed in our studies.

Importantly, our results showing that SUMO2 inhibition is effective in cells driven by different SySa fusions, irrespective of their carboxy-terminal fusion partner (SSX1 or SSX2) indicate that these inhibitors may be broadly applicable for SySa patients with distinct SySa fusion proteins. Also, since targeted therapies are more likely to be successful in combination with other cytotoxic agents, SUMO2 inhibitors may work more effectively in combination with currently used chemotherapies. Taken together, our results highlight the potential of SUMO2 inhibitors as promising therapeutic targets for SySa, with TAK-981 emerging as a particularly strong candidate for clinical testing in patients with synovial sarcoma.

## METHODS

### DepMap data mining and library construction

To identify SS-specific dependencies, we filtered the DepMap CRISPR as well as the RNAi Achilles dataset for genes that were more dependent on growth for synovial sarcoma cell lines when compared to all other cancer cell lines. We then selected the top 200 genes from each dataset which resulted in 348 unique genes in the combined dataset. sgRNA for these genes were designed using CRISPick tool^36^ from the Broad Institute. sgRNA libraries were synthesized using Array technology (CustomArray, Inc.) containing 3665 guides targeting 348 genes along with 174 guides as non-targeting controls. The guides were amplified by PCR and cloned into pKLO.1 by ligation using the Esp3I (NEB) restriction sites^39,40^. Transformations were performed with Invitrogen’s MegaX DH10B T1 electro-competent cells using an Eppendorf electroporator 2510 and Bio-Rad Gene Pulser 1 mm cuvettes. A minimum of 30 million successfully transformed cells or 30,000X coverage of the library was obtained.

### Cell culture

HEK-293T and SYO1 cells were cultured in DMEM supplemented with 10% FBS, 1 % penicillin-streptomycin and 1% L-glutamine. HS-SY-II and HS-SY-II-Cas9 cells were additionally supplemented with 0.5% Sodium Pyruvate. Aska-SS and Yamato-SS cells were maintained in DMEM supplemented with 20% FBS, 1% penicillin-streptomycin and 1% L-glutamine. The 1273/99 cell line was cultured in DMEM supplemented with F12. All cell lines were authenticated by STR profiling.

### Virus production

Lentivirus was produced in HEK293T cells. Cells from four 80% confluent 10 cm Petri dishes were transfected with 0.9μg VSV-G envelope expressing plasmid pMD2 and 9 μg psPAX2 packaging vectors and 9μg of the gRNA library DNA in the presence of 113.4μL Polyethylenimine - PEI (VWR International, 1 mg/mL) per plate. Medium was exchanged after overnight incubation and virus supernatant was collected after 48 and 72 hours, passed through a 0.45 μm filter and concentrated by centrifuging at 6,000 g for 2 hours at 4°C. Supernatant was discarded, and pellets were resuspended in 1/1,000^th^ volume of PBS and rotated at 4°C overnight. The concentrated virus was flash frozen in ethanol-dry ice bath and stored at −80°C.

### *In vitro* and *in vivo* CRISPR/Cas9 screens

Screens were performed in duplicates. HS-SY-II Cas9 cells were transduced with the screen library in the presence of 0.8 mg/ml polybrene with an efficiency of 30% or lower to ensure most cells received a single sgRNA. After selection with puromycin (1 μg/mL) for 2-4 days, a cell aliquot containing 5 million cells (∼1,000X coverage of library) was frozen as the day 0 or input reference sample. The remaining cells were divided into 2 arms for the *in vivo* and *in vitro* screens. For the *in vivo* screen, 2 million cells in 50% Matrigel were transplanted subcutaneously into the flanks of 4 athymic nude mice per replicate. The resultant tumor was monitored, and mice were sacrificed when the tumor volume reached 1 cm^3^. The tumor was dissociated into single cell suspension^41^ using collagenase II (20 mg/mL) along with Dnase I (10,000 Kunitz/mL) and used for further experiments. For the *in vitro* screens, at least 5 million cells were maintained throughout the 14-day culture period and collected at the end of the screen. Genomic DNA was extracted from collected cell pellets using a Zymo Quick DNA miniprep kit (#D3024). The sgRNA were PCR amplified by NEBNext Ultra II Q5 Master Mix (NEB #M0544) from the genomic DNA using the indexed PCR primers with next-generation sequencing adapters compatible with Illumina’s NEXTERA kit. PCR products were size-selected by gel electrophoresis, quantified by Qubit (Thermo Fisher Scientific) and sequenced using HiSeq (Illumina).

### Proliferation assay

The proliferation assay was performed using CellTiter-Glo Luminescent Cell Viability Assay (Promega #G7570) using manufacturer’s instructions. Cell numbers were optimized for 384-well plate for each cell line. SUMO2 inhibitor TAK-981 dissolved in DMSO were echo dotted on to a 384-well plate in varying concentrations with the final concentration of DMSO at 0.08% in each well. 25ul of 50,000 cells/ml were seeded in each well of a 384-well plate. The cells were incubated at 37°C at 5% CO_2_ for 48 hours then quenched with CellTiter-Glo^®^, centrifuged at 1000 rpm for 1 min and incubated at RT for 20 min. Luminescence was recorded with a plate reader (BMG FLUOStar). EC_50_ values were calculated by GraphPad Prism software.

### Colony forming assays

Crystal violet colony-forming assays were conducted by seeding cells at low density in a 6-well plate. After the cells adhered to the plate, they were treated to varying concentrations of TAK-981 for an additional 2-4 days. The wells were then washed, fixed and stained with 0.02% crystal violet solution in methanol. Subsequently, wells were imaged for quantification.

### 2D and 3D cell culture and TAK-981 treatment

To culture cells in 2D and 3D growth formats, SYO1 and ASKA-SS cells were grown in high-glucose DMEM supplemented with 1% L-glutamine (Gibco #11965092) and 1% antibiotic-antimycotic (Gibco #15240062) and supplemented with 10% and 20% FBS (Gibco #16140071) respectively. Cells were trypsinized (Gibco #25300054) and counted using Cellometer Auto 2000 (Nexcelom). Mammocult medium (StemCell Technologies #50620) with the addition of 0.5% Hydrocortisone (StemCell Technologies #07925) and 0.2% Heparin (StemCell Technologies #07980) was used as described previously^42^.

For the 2D experiments, cells were resuspended in Mammocult at a concentration of 50,000 cells/ml. 100 μl of the solution was dispensed in each well of a 96-well plate. For 3D experiments, cells were resuspended at a concentration of 500,000 cells/ml in a 3:4 mixture solution of Mammocult medium and Matrigel (Corning #354234). The mixture was kept on ice throughout the seeding process. 10 μl of this solution was dispensed around the perimeter of each well’s bottom of a 96-well plate to create mini-rings as established previously^42–45^. After a 30-minute incubation at 37°C to solidify the gel, 100 μl of pre-warmed Mammocult medium to was added to each well using an automated fluid handler (Microlab NIMBUS, Hamilton). In all cases, plates were imaged in brightfield mode every 24 hours using a high-content microscope (Celigo, Nexcelom).

Plates were incubated for 2 days before initiating drug treatment. Pre-warmed Mammocult medium containing TAK-981 (MedChemExpress #HY-111789) at six different concentrations diluted in DMSO (Fisher Scientific #BP231-100) was added to the plates after complete removal of media. Each plate included 10 μM staurosporine (Selleckchem #S1421) and 1% DMSO as positive and negative controls respectively.

Treatment was repeated twice or three times after subsequent 24-hour incubations. Cell viability was measured after 2 days (post-two total treatments), 3 or 4 days (post-three total treatments) of incubation with TAK-981. Viability was assessed via ATP-release assay (CellTiter-Glo 3D, Promega #PRG9683) after PBS washes (Gibco #14190144) and incubation with 50 μl of dispase (Gibco #17105041) at 37°C for 25 minutes. Plates were incubated in the dark at room temperature for 25 minutes upon addition of the CellTiter Glo 3D reagent. Luminescence was measured using a SpectraMax iD3 plate reader (Molecular Devices). The viability of each well was normalized to the vehicle control wells.

### Apoptosis and cell cycle assays

Apoptosis was quantified by flow cytometry using Annexin V-FITC kit from BD Biosciences. 3*10^5^ SYO1 and HS-SY-II cells were seeded in a 6-well plate and allowed to attach for 24 hours. TAK-981 was added in varying concentrations and incubated for 48 hours. After incubation, the cells were trypsinized, washed in warm PBS, and resuspended in Annexin V binding buffer. Annexin V-FITC was added and incubated at room temperature for 10 minutes. The samples were then analyzed by flow cytometry using Fortessa (BD Bioscience, USA) along with FlowJo analysis software.

### Cell cycle analysis

Cell cycle analysis was done by staining the cells with propidium iodide (PI). As previously stated, 3*10^5^ SYO1 and HS-SY-II cells were seeded in a 6-well plate and allowed to attach for 24 hours. They were then exposed to varying concentrations of TAK-981 for 48 hours. Cells were trypsinized, washed with PBS, and fixed with ethanol. Cells were washed and stained with PI. The samples were then analyzed by flow cytometry using Fortessa (BD Bioscience, USA) along with FlowJo analysis software.

### RNA sequencing

HS-SY-II and SYO1 cells were treated with either TAK-981 at concentrations of 25nM and 100nM respectively for the treatment arm or DMSO for the control arm for 48 hours. Cells were pelleted and RNA was extracted using Trizol (Thermo, Cat.No. 15596026) with concentration determined by Qubit (Thermo Scientific). Libraries were prepared with the NEBNext Ultra II RNALibrary prep kit for Illumina (NEB, Cat.No. E7770S).

### Small hairpin RNA (shRNA) transfection and transduction

Small hairpin RNA (shRNA) for SUMO2 were cloned into the all-in-one-Tet vector and packaged into lentivirus using pMD and pPax2 as described above. 300,000 HS-SY-II and SYO1 cells were seeded into six-well culture plates overnight. Lipofectamine 2000 reagent (Invitrogen, Waltham, MA, USA) was used to perform the transfections, as described in the manufacturer’s instructions. At 48 h after transfection, media was changed and puromycin selected for 2 days. After selection, cells were subjected to a previously determined amount of doxycycline (4.5ug/ml) for 48 hours. Cells were harvested for qPCR quantification and western blot analysis.

### Quantitative real-time reverse transcription PCR (qRT-PCR)

TRIzol Reagent (Thermo Fisher) was used to extract total RNA from SYO1 and HS-SY-II cell pellets and 1^st^ strand was synthesized using Protoscript II (NEB) with polyA selection. qPCR was performed using TaqMan Gene Expression Master Mix and FAM probes for SUMO2, HPRT, GAPDH from Thermo Fisher.

### Western blot analysis

Whole cell lysates from synovial sarcoma cancer cell lines treated with varying concentrations of TAK-981 or with 0.1% DMSO were prepared on ice with RIPA lysis buffer (Thermo #89900) supplemented with protease inhibitor cocktail (Thermo #78429). Lysates along with LDS Sample buffer (Thermo #J61942.AD) were heated at 65 °C for 10 min. Proteins were separated on precast 4%–12% Bis-Tris gradient gels (Thermo #NW04120BOX). Separated proteins were subsequently transferred to nitrocellulose membranes (Thermo #IB23001) using the iBLOT2 system. Membranes were blocked with PBS containing 5% milk powder and 0.05% Tween-20 for 1 hour. Protein samples were incubated with primary antibodies against SS18-SSX fusion at 1:1000 dilution (rabbit monoclonal from Cell Signaling Technologies #70929), SUMO2/3 at 1:500 dilution (mouse monoclonal 8A2 from Abcam #ab81371) and H2AK119ub at 1:1000 dilution (rabbit monoclonal from Cell Signaling Technologies #8240). Mouse anti-Vinculin monoclonal antibody (Abcam #ab130007) was used as a loading control. (Goat anti-rabbit IgG-HRP and goat anti-mouse IgG-HRP were used as secondary antibodies at 1:3000 dilution in 5% milk. Signal was detected using SuperSignal West Femto (Thermo # 34094) and captured using the BioRad ChemiDoc system (Cat.No.1708370).

### *In vivo* tumor models

In this study, 8- to 10-week-old male Nu/J mice were acquired from Jackson Laboratory. The animals were housed in individually ventilated cages under specified pathogen-free conditions in the animal facilities of our institute.

Two million Aska-SS or SYO1 cells were injected subcutaneously into the right flank of each mouse in a mixture of 100 µL PBS and 50% Matrigel (Corning #356234). Once tumors were established, typically within 3-8 days post-implantation, drug treatment was initiated. Mice received 0.25 mL of 25 mg/kg TAK-981 in 20% HPBCD or a vehicle control via intraperitoneal injection three times per week. Tumor growth was monitored two to three times per week using a vernier caliper and imaging until ethical endpoints necessitated euthanasia due to tumor size or ulceration. Tumor volume was calculated using the formula: volume (V) = W² × L / 2, where W is the width and L is the length of the tumor.

### Immunohistochemistry

Tumors excised from nude mice were formalin-fixed and paraffin-embedded. Sections of 5 μm were cut and mounted on slides (Medline Cat.No. MLABSLIDE1WC). After deparaffinization, antigen retrieval was carried out in PBS (pH 6) using a pressure cooker for 10-15 minutes. Tissue sections were blocked with 10% donkey serum for an hour and incubated with the primary antibody at 4C overnight. After multiple PBS washes, the sections were incubated with the secondary antibody for 45 minutes at room temperature. Visualization was performed using HRP substrate DAB (3, 3 -diaminobenzidine) (Cat. No. SK-4105). Sections were counterstained with hematoxylin.

### ChIP-seq

SYO1 cells were treated with DMSO (control) or 1 uM TAK-981 for 72hrs. ChIP-seq was performed to assess changes in histone 2A ubiquitination at lysine 119 (H2AK119ub) as described earlier^46^. SYO1 cells plated and treated in 10 cm tissue culture treated plates in triplicates for each group (DMSO and TAK-981) were trypsinized and counted for fixing after 72 hr treatment. 1 million cells from each plate were fixed using 1% formaldehyde for 10 minutes at room temperature. Fixed cells were sheared using Bioruptor (Diagenode, NJ) in 15 cycles, each with 30 sec. on and 30 sec. off settings at 4^0^C. Chromatin was immunoprecipitated using antibody for H2AK119ub (Cell Signaling Technologies # 8240). DNA was purified after reverse crosslinking. Immunoprecipitated chromatin was subjected to library prep using NEBNext Ultra II DNA library prep kit for Illumina (E7645S and E7600S) as per the manufacturer’s protocol. Library prepped DNA then sequenced on AVITI platform (Element Biosciences) with the 2×75bp High Output Cloudbreak Freestyle Kit.

### CUT&RUN

Changes in genome wide binding of the SS18-SSX2 fusion after TAK-981 treatment were studied in SYO1 cell line using CUT&RUN assay. SYO1 cells were treated in duplicates with DMSO or 1 uM TAK-981 for 72 hr. At 72 hrs, cells were trypsinized, washed with PBS and counted for the assay. 300,000 cells per antibody were then bound on activated ConA magnetic beads and CUT&RUN was performed using the CUTANA ChIC/CUT&RUN kit (Epicypher, NC # 14-1048) as per the manufacturer’s protocol. Permeabilized cells were incubated with anti-Rabbit IgG (Epicypher, #13-0042) or SSX-SS18 (Cell Signaling technologies # 72364) overnight at 4^0^C. K-MetStat panel (provided in the kit, #19-1002) was added to IgG control samples. E.coli Spike-In DNA, also provided in the kit (#18-1401) was added to each sample as mentioned in the protocol. Purified DNA then subjected to library prep using NEBNext Ultra II DNA library prep kit for Illumina (E7645S and E7600S) and sequenced on Element Biosciences AVITI platform with the 2×75bp High Output Cloudbreak Freestyle Kit.

### Data Analysis

#### Pooled CRISPR screen

MAGeCK (Model-based Analysis of Genome-wide CRISPR-Cas9 Knockout)^28^ pipeline was used for mapping reads (paired end fastqs) to sgRNA custom library (Supplemental Table S1) from DepMap (DepMap repository version 21Q3 https://depmap.org/portal/), normalization using 10% of non-targeting controls (Supplemental Table S1), and quality control. Identification of positively and negatively selected genes/hits comparing Day14_in_vitro vs Day0 and CDX vs Day0.was performed used MAGeCK RRA (Robust Rank Aggregation).

#### CUT&RUN

Paired-end reads were trimmed using Cutadapt version 2.3^47^ with parameters “-j 12 -m 20 -O 5 - q 15 -a AGATCGGAAGAGCACACGTCTGAACTCCAGTCAC -A AGATCGGAAGAGCGTCGTGTAGGGAAAGAGTGT”. Trimmed reads were aligned against E. coli genomic sequence (GCF_000005845.2_ASM584v2_genomic) using Bowtie2 version 2.2.5^48^ with parameters “--local” to quantify spike-in amount. Unmapped reads were subsequently aligned against hg38 chrM to remove mitochondrial reads using Bowtie2 with parameters “—local -X 2000”. Remaining unaligned reads were mapped against human genome version hg38 (without chrM) using Bowtie2 with parameters “--very-sensitive --no-discordant -X 2000”. Multimapping and improperly paired reads were removed using Deeptools alignmentSieve version 3.4.3^49^ with parameters “--minMappingQuality 30 --samFlagInclude 2”. Duplicate reads were removed using Picard MarkDuplicates version 2.22.0. Peak calling was performed in DMSO treated samples using Macs2 version 2.2.9.1^50^ with parameters “--nomodel --shift -75 --extsize 150 --keep-dup all -q 0.01 --broad --broad-cutoff 0.1 --gsize 2700000000.0 --format BAMPE”. Overlapping peaks in DMSO replicates were determined using Bedtools intersect version 2.29.2^51^. Overlapping peaks were merged using Bedtools merge and parameter “-d 500”. Peaks overlapping Encode blacklist regions were removed. Peaks were annotated and peak tags were counted in each sample using Homer annotatePeaks.pl^52^ with parameters “hg38 -raw”. Bigwig signal files were generated using Deeptools bamCoverage with parameters “--binSize 20 -- smoothLength 500 -p 12 --normalizeUsing RPGC --extendReads --ignoreForNormalization chrX--effectiveGenomeSize 2913022398 –scaleFactor [*spike-in scale factor*]”.

#### ChIP-seq

Mouse spike-in reads were classified and separated from SYO1 ChIP-seq reads using Xenome version 1.0.0^53^. First mate (read1) of each sample were trimmed using Cutadapt version 2.3^47^ with parameters “-j 12 -m 30 -O 5 -q 15 -a AGATCGGAAGAGCACACGTCTGAACTCCAGTCAC”. Trimmed reads were aligned against hg38 chrM to remove mitochondrial reads using Bowtie2^48^ with parameters “--local”. Remaining unaligned reads were mapped against human genome version hg38 (without chrM) using Bowtie2 with parameters “--local”. Multimapping reads were removed using Deeptools alignment Sieve version 3.4.3^49^ with parameters “--minMappingQuality 30”. Duplicate reads were removed using Picard MarkDuplicates version 2.22.0. Peak calling was performed using Macs2 version 2.2.9.1^50^ with parameters “--keep-dup all -q 0.01 --broad --broad-cutoff 0.1 --gsize 2700000000.0 --format BAM”. Overlapping peaks in DMSO replicates were determined using Bedtools intersect version 2.29.2^51^. Overlapping peaks were merged using Bedtools merge. Peaks overlapping Encode blacklist regions were removed. Peaks were annotated and peak tags were counted in each sample using Homer annotatePeaks.pl^52^ with parameters “hg38 -raw”. Bigwig signal files were generated using Deeptools bamCoverage with parameters “--binSize 20 --smoothLength 500 -p 12 --normalizeUsing RPGC -- ignoreForNormalization chrX --effectiveGenomeSize 2913022398

#### RNA-seq data analysis

Raw reads were preprocessed by trimming Illumina Truseq adapters, polyA, and polyT sequences using cutadapt v2.313 with parameters “cutadapt -j 4 -m 20 --interleaved -a AGATCGGAAGAGCACACGTCTGAACTCCAGTCAC -A AGATCGGAAGAGCGTCGTGTAGGGAAAGAGTGT Fastq1 Fastq2 | cutadapt --interleaved -j 4-m 20 -a “A{100}” -A “A{100}” - | cutadapt -j 4 -m 20 -a “T{100}” -A “T{100}” -”. Trimmed reads were subsequently aligned to human genome version hg38 using STAR aligner v2.7.0d_0221 14 with parameters according to ENCODE long RNA-seq pipeline (https://github.com/ENCODE-DCC/long-rna-seq-pipeline). Gene expression levels were quantified using RSEM v1.3.1 15. Ensembl v84 gene annotations were used for the alignment and quantification steps. RNA-seq sequence, alignment, and quantification qualities were assessed using FastQC v0.11.5 (https://www.bioinformatics.babraham.ac.uk/projects/fastqc/) and MultiQC v1.8 16. Lowly expressed genes were filtered out by retaining genes with estimated counts (from RSEM) ≥ number of samples times 5. Filtered estimated read counts from RSEM were used for differential expression comparisons using the Wald test implemented in the R Bioconductor package DESeq2 v1.22.2 based on generalized linear model and negative binomial distribution 17^54^. Genes with Benjamini-Hochberg corrected p-value < 0.05 and fold change ≥ 2.0 or ≤ 2.0 were selected as differentially expressed genes. Gene set enrichment analysis (GSEA) was performed using GSEA app version 4.3.2^55^.

#### Data availability

Sequencing data for RNA-seq, CUT&RUN, ChIP-seq, and high-throughput CRISPR screens, are deposited in the NCBI GEO under accession number: GSE276074. Code for data analysis is available in the Github page : https://github.com/PBioinfo/Synovial_Sarcoma_Paper_code.git

### Statistical analysis

All statistical analysis was performed using GraphPad Prism 9.0. Data were presented as the mean ± SEM. Statistical significance between 2 groups was determined using Students t-test. Significance over multiple time points among groups was computed using 2-way ANOVA. Dose response curves were fit using four parameter logistic equation. A statistical threshold of *P* < 0.05 was used with ^∗^,*P* < 0.05; ^∗∗^, *P* < 0.01; ^∗∗∗^, *P* < 0.001; \*\*\*\**P* < 0.0001; *P* =ns, not significant.

## ACKNOWLEDGEMENTS

We would like to thank Adriana Charbono and Buddy Charbono for their invaluable assistance with mouse studies, Dr. Chih-Cheng Yang and Chun-Teng Huang from the Sanford Burnham Prebys Medical Discovery Institute (SBP) functional genomics core, Yoav Altman from the SBP Flow Cytometry Core, and Drs. Rebecca Porritt and Kang Liu from the Genomics Core for their excellent support. We would like to acknowledge the help of Dr. Derron Herr and Anis Shahnaee from the Jerold Chun Lab at SBP for their assistance with microscopy. Some schematic figures were made using Biorender.com.

This work was supported by National Institutes of Health (NIH) National Cancer Institute grants CA262746 and P30 CA030199. We would also like to acknowledge the support of the Animal Facility and the SBP Flow Cytometry Core supported by the NCI Cancer Center Support Grant P30 CA030199.

## AUTHOR CONTRIBUTIONS

**RI.:** conceptualization, data curation, formal analysis, investigation, methodology, project administration, validation, visualization, writing-original draft. **AD**.: conceptualization, data curation, formal analysis**, AP.:** conceptualization, data curation, formal analysis**, TK.:** conceptualization, data curation, formal analysis**, GB.:** resources, **SAA.**: resources, **D.F.:** data curation, formal analysis, methodology, **R.M**: data curation, formal analysis, methodology, software. **P.A.B.:** data curation, formal analysis, methodology, software. **K.V.**: resources, funding acquisition, **AS.**; conceptualization, resources, funding acquisition, **A.J.D.:** conceptualization, resources, funding acquisition, investigation, methodology, project administration, supervision, visualization, writing-original draft.

## Conflict-of-interest disclosure

The authors declare no competing interests or conflicts of interest related to this work.

## Notes

### Competing Interest Statement

The authors have declared no competing interest.

## REFERENCES

1. Mastrangelo, G., Coindre, J.-M., Ducimetière, F., Dei Tos, A.P., Fadda, E., Blay, J.-Y., Buja, A., Fedeli, U., Cegolon, L., Frasson, A., et al. (2012). Incidence of soft tissue sarcoma and beyond: a population-based prospective study in 3 European regions. Cancer 118, 5339–5348.

2. Moch, H. (2020). Soft Tissue and Bone Tumours WHO Classification of Tumours / Volume 3. In WHO Classification of Tumours WHO Classification of Tumours., H. Moch, ed. (International Agency for Research on Cancer).

3. Speth, B.M., Krieg, A.H., Kaelin, A., Exner, G.U., Guillou, L., von Hochstetter, A., Jundt, G., and Hefti, F. (2011). Synovial sarcoma in patients under 20 years of age: a multicenter study with a minimum follow-up of 10 years. J. Child. Orthop. 5, 335–342.

4. Sultan, I., Rodriguez-Galindo, C., Saab, R., Yasir, S., Casanova, M., and Ferrari, A. (2009). Comparing children and adults with synovial sarcoma in the Surveillance, Epidemiology, and End Results program, 1983 to 2005: an analysis of 1268 patients. Cancer 115, 3537–3547.

5. Ladanyi, M. (1995). The emerging molecular genetics of sarcoma translocations. Diagn. Mol. Pathol. 4, 162–173.

6. Sorensen, P., and Triche, T.J. (1996). Gene fusions encoding chimeric transcription factors in solid tumors. Semin Cancer Biol 7, 3–14.

7. McBride, M.J., Pulice, J.L., Beird, H.C., Ingram, D.R., D’Avino, A.R., Shern, J.F., Charville, G.W., Hornick, J.L., Nakayama, R.T., Garcia-Rivera, E.M., et al. (2018). The SS18-SSX fusion oncoprotein hijacks BAF complex targeting and function to drive synovial sarcoma. Cancer Cell 33, 1128–1141.e7.

8. Weber, C.M., Hafner, A., Kirkland, J.G., Braun, S.M.G., Stanton, B.Z., Boettiger, A.N., and Crabtree, G.R. (2021). mSWI/SNF promotes Polycomb repression both directly and through genome-wide redistribution. Nat. Struct. Mol. Biol. 28, 501–511.

9. Barham, W., Frump, A.L., Sherrill, T.P., Garcia, C.B., Saito-Diaz, K., VanSaun, M.N., Fingleton, B., Gleaves, L., Orton, D., Capecchi, M.R., et al. (2013). Targeting the Wnt pathway in synovial sarcoma models. Cancer Discov. 3, 1286–1301.

10. Ng, T.L., Gown, A.M., Barry, T.S., Cheang, M.C.U., Chan, A.K.W., Turbin, D.A., Hsu, F.D., West, R.B., and Nielsen, T.O. (2005). Nuclear beta-catenin in mesenchymal tumors. Mod. Pathol. 18, 68–74.

11. DeSalvo, J., Ban, Y., Li, L., Sun, X., Jiang, Z., Kerr, D.A., Khanlari, M., Boulina, M., Capecchi, M.R., Partanen, J.M., et al. (2021). ETV4 and ETV5 drive synovial sarcoma through cell cycle and DUX4 embryonic pathway control. J. Clin. Invest. 131. 10.1172/JCI141908.

12. Rota, R., Ciarapica, R., Miele, L., and Locatelli, F. (2012). Notch signaling in pediatric soft tissue sarcomas. BMC Med. 10, 141.

13. Ciarapica, R., Miele, L., Giordano, A., Locatelli, F., and Rota, R. (2011). Enhancer of zeste homolog 2 (EZH2) in pediatric soft tissue sarcomas: first implications. BMC Med. 9, 63.

14. Su, L., Cheng, H., Sampaio, A.V., Nielsen, T.O., and Underhill, T.M. (2010). EGR1 reactivation by histone deacetylase inhibitors promotes synovial sarcoma cell death through the PTEN tumor suppressor. Oncogene 29, 4352–4361.

15. CDKN2A) Gene Deletion Is a Frequent Genetic Event in Synovial Sarcomas 16I-K20.

16. McBride, M.J., Mashtalir, N., Winter, E.B., Dao, H.T., Filipovski, M., D’Avino, A.R., Seo, H.-S., Umbreit, N.T., St Pierre, R., Valencia, A.M., et al. (2021). Author Correction: The nucleosome acidic patch and H2A ubiquitination underlie mSWI/SNF recruitment in synovial sarcoma. Nat. Struct. Mol. Biol. 28, 118.

17. Benabdallah, N.S., Dalal, V., Scott, R.W., Marcous, F., Sotiriou, A., Kommoss, F.K.F., Pejkovska, A., Gaspar, L., Wagner, L., Sánchez-Rivera, F.J., et al. (2023). Aberrant gene activation in synovial sarcoma relies on SSX specificity and increased PRC1.1 stability. Nat. Struct. Mol. Biol. 30, 1640–1652.

18. Hart, T., Chandrashekhar, M., Aregger, M., Steinhart, Z., Brown, K.R., MacLeod, G., Mis, M., Zimmermann, M., Fradet-Turcotte, A., Sun, S., et al. (2015). High-resolution CRISPR screens reveal fitness genes and genotype-specific cancer liabilities. Cell 163, 1515–1526.

19. Dempster, J.M., Pacini, C., Pantel, S., Behan, F.M., Green, T., Krill-Burger, J., Beaver, C.M., Younger, S.T., Zhivich, V., Najgebauer, H., et al. (2019). Agreement between two large pan-cancer CRISPR-Cas9 gene dependency datasets. bioRxiv. 10.1101/604447.

20. GTEx Consortium (2020). The GTEx Consortium atlas of genetic regulatory effects across human tissues. Science 369, 1318–1330.

21. Doench, J.G., Hartenian, E., Graham, D.B., Tothova, Z., Hegde, M., Smith, I., Sullender, M., Ebert, B.L., Xavier, R.J., and Root, D.E. (2014). Rational design of highly active sgRNAs for CRISPR-Cas9-mediated gene inactivation. Nat. Biotechnol. 32, 1262–1267.

22. Jung, H.R., Oh, Y., Na, D., Min, S., Kang, J., Jang, D., Shin, S., Kim, J., Lee, S.E., Jeong, E.M., et al. (2021). CRISPR screens identify a novel combination treatment targeting BCL-XL and WNT signaling for KRAS/BRAF-mutated colorectal cancers. Oncogene 40, 3287–3302.

23. Behan, F.M., Iorio, F., Picco, G., Gonçalves, E., Beaver, C.M., Migliardi, G., Santos, R., Rao, Y., Sassi, F., Pinnelli, M., et al. (2019). Prioritization of cancer therapeutic targets using CRISPR-Cas9 screens. Nature 568, 511–516.

24. Brien, G.L., Remillard, D., Shi, J., Hemming, M.L., Chabon, J., Wynne, K., Dillon, E.T., Cagney, G., Van Mierlo, G., Baltissen, M.P., et al. (2018). Targeted degradation of BRD9 reverses oncogenic gene expression in synovial sarcoma. Elife 7. 10.7554/eLife.41305.

25. Chen, E.Y., Tan, C.M., Kou, Y., Duan, Q., Wang, Z., Meirelles, G.V., Clark, N.R., and Ma’ayan, A. (2013). Enrichr: interactive and collaborative HTML5 gene list enrichment analysis tool. BMC Bioinformatics 14, 128.

26. Michel, B.C., D’Avino, A.R., Cassel, S.H., Mashtalir, N., McKenzie, Z.M., McBride, M.J., Valencia, A.M., Zhou, Q., Bocker, M., Soares, L.M.M., et al. (2018). A non-canonical SWI/SNF complex is a synthetic lethal target in cancers driven by BAF complex perturbation. Nat. Cell Biol. 20, 1410–1420.

27. Conant, D., Hsiau, T., Rossi, N., Oki, J., Maures, T., Waite, K., Yang, J., Joshi, S., Kelso, R., Holden, K., et al. (2022). Inference of CRISPR edits from Sanger trace data. CRISPR J. 5, 123–130.

28. Li, W., Xu, H., Xiao, T., Cong, L., Love, M.I., Zhang, F., Irizarry, R.A., Liu, J.S., Brown, M., and Liu, X.S. (2014). MAGeCK enables robust identification of essential genes from genome-scale CRISPR/Cas9 knockout screens. Genome Biol. 15, 554.

29. Jerby-Arnon, L., Neftel, C., Shore, M.E., Weisman, H.R., Mathewson, N.D., McBride, M.J., Haas, B., Izar, B., Volorio, A., Boulay, G., et al. (2021). Opposing immune and genetic mechanisms shape oncogenic programs in synovial sarcoma. Nat. Med. 27, 289–300.

30. Barreto Coelho, P., Costa, P.A., Espejo Freire, A.P., Kwon, D., Jonczak, E., D’Amato, G.Z., and Trent, J.C. (2021). Outcomes of metastatic synovial sarcoma with doxorubicin, pazopanib, and ifosfamide therapy. J. Clin. Oncol. 39, e23552–e23552.

31. Kadoch, C., and Crabtree, G.R. (2013). Reversible disruption of mSWI/SNF (BAF) complexes by the SS18-SSX oncogenic fusion in synovial sarcoma. Cell 153, 71–85.

32. Boulay, G., Cironi, L., Garcia, S.P., Rengarajan, S., Xing, Y.-H., Lee, L., Awad, M.E., Naigles, B., Iyer, S., Broye, L.C., et al. (2021). The chromatin landscape of primary synovial sarcoma organoids is linked to specific epigenetic mechanisms and dependencies. Life Sci Alliance 4. 10.26508/lsa.202000808.

33. Tong, Z., Ai, H., Xu, Z., He, K., Chu, G.-C., Shi, Q., Deng, Z., Xue, Q., Sun, M., Du, Y., et al. (2024). Synovial sarcoma X breakpoint 1 protein uses a cryptic groove to selectively recognize H2AK119Ub nucleosomes. Nat. Struct. Mol. Biol. 31, 300–310.

34. McBride, M.J., Mashtalir, N., Winter, E.B., Dao, H.T., Filipovski, M., D’Avino, A.R., Seo, H.-S., Umbreit, N.T., St Pierre, R., Valencia, A.M., et al. (2020). The nucleosome acidic patch and H2A ubiquitination underlie mSWI/SNF recruitment in synovial sarcoma. Nat. Struct. Mol. Biol. 27, 836–845.

35. Kim, H.S., Kim, B.-R., Dao, T.T.P., Kim, J.-M., Kim, Y.-J., Son, H., Jo, S., Kim, D., Kim, J., Suh, Y.J., et al. (2023). TAK-981, a SUMOylation inhibitor, suppresses AML growth immune-independently. Blood Adv. 7, 3155–3168.

36. Doench, J.G., Fusi, N., Sullender, M., Hegde, M., Vaimberg, E.W., Donovan, K.F., Smith, I., Tothova, Z., Wilen, C., Orchard, R., et al. (2016). Optimized sgRNA design to maximize activity and minimize off-target effects of CRISPR-Cas9. Nat. Biotechnol. 34, 184–191.

37. Kumar, S., Schoonderwoerd, M.J.A., Kroonen, J.S., de Graaf, I.J., Sluijter, M., Ruano, D., González-Prieto, R., Verlaan-de Vries, M., Rip, J., Arens, R., et al. (2022). Targeting pancreatic cancer by TAK-981: a SUMOylation inhibitor that activates the immune system and blocks cancer cell cycle progression in a preclinical model. Gut 71, 2266–2283.

38. Lightcap, E.S., Yu, P., Grossman, S., Song, K., Khattar, M., Xega, K., He, X., Gavin, J.M., Imaichi, H., Garnsey, J.J., et al. (2021). A small-molecule SUMOylation inhibitor activates antitumor immune responses and potentiates immune therapies in preclinical models. Sci. Transl. Med. 13, eaba7791.

39. Wang, T., Lander, E.S., and Sabatini, D.M. (2016). Large-scale single guide RNA library construction and use for CRISPR-Cas9-based genetic screens. Cold Spring Harb. Protoc. 2016, db.top086892.

40. Wang, T., Lander, E.S., and Sabatini, D.M. (2016). Single guide RNA library design and construction. Cold Spring Harb. Protoc. 2016, db.prot090803.

41. Preparing Viable Single Cells from Human Tissue and Tumors for Cytomic Analysis Current Protocols in Molecular Biology UNIT 25C.1 Biology UNIT 25.

42. Al Shihabi, A., Davarifar, A., Nguyen, H.T.L., Tavanaie, N., Nelson, S.D., Yanagawa, J., Federman, N., Bernthal, N., Hornicek, F., and Soragni, A. (2022). Personalized chordoma organoids for drug discovery studies. Sci. Adv. 8, eabl3674.

43. Phan, N., Hong, J.J., Tofig, B., Mapua, M., Elashoff, D., Moatamed, N.A., Huang, J., Memarzadeh, S., Damoiseaux, R., and Soragni, A. (2019). A simple high-throughput approach identifies actionable drug sensitivities in patient-derived tumor organoids. Commun. Biol. 2, 78.

44. Nguyen, H.T.L., and Soragni, A. (2020). Patient-derived tumor organoid rings for histologic characterization and high-throughput screening. STAR Protoc. 1, 100056.

45. Tebon, P.J., Wang, B., Markowitz, A.L., Davarifar, A., Tsai, B.L., Krawczuk, P., Gonzalez, A.E., Sartini, S., Murray, G.F., Nguyen, H.T.L., et al. (2023). Drug screening at single-organoid resolution via bioprinting and interferometry. Nat. Commun. 14, 3168.

46. Barbosa, K., Deshpande, A., Perales, M., Xiang, P., Murad, R., Pramod, A.B., Minkina, A., Robertson, N., Schischlik, F., Lei, X., et al. (2024). Transcriptional control of leukemogenesis by the chromatin reader SGF29. Blood 143, 697–712.

47. Martin, M. (2011). Cutadapt removes adapter sequences from high-throughput sequencing reads. EMBnet J. 17, 10.

48. Langmead, B., and Salzberg, S.L. (2012). Fast gapped-read alignment with Bowtie 2. Nat. Methods 9, 357–359.

49. Ramírez, F., Dündar, F., Diehl, S., Grüning, B.A., and Manke, T. (2014). deepTools: a flexible platform for exploring deep-sequencing data. Nucleic Acids Res. 42, W187–91.

50. Zhang, Y., Liu, T., Meyer, C.A., Eeckhoute, J., Johnson, D.S., Bernstein, B.E., Nusbaum, C., Myers, R.M., Brown, M., Li, W., et al. (2008). Model-based analysis of ChIP-Seq (MACS). Genome Biol. 9, R137.

51. Quinlan, A.R., and Hall, I.M. (2010). BEDTools: a flexible suite of utilities for comparing genomic features. Bioinformatics 26, 841–842.

52. Heinz, S., Benner, C., Spann, N., Bertolino, E., Lin, Y.C., Laslo, P., Cheng, J.X., Murre, C., Singh, H., and Glass, C.K. (2010). Simple combinations of lineage-determining transcription factors prime cis-regulatory elements required for macrophage and B cell identities. Mol. Cell 38, 576–589.

53. Conway, T., Wazny, J., Bromage, A., Tymms, M., Sooraj, D., Williams, E.D., and Beresford-Smith, B. (2012). Xenome--a tool for classifying reads from xenograft samples. Bioinformatics 28, i172–8.

54. Love, M.I., Huber, W., and Anders, S. (2014). Moderated estimation of fold change and dispersion for RNA-seq data with DESeq2. bioRxiv. 10.1101/002832.

55. Subramanian, A., Tamayo, P., Mootha, V.K., Mukherjee, S., Ebert, B.L., Gillette, M.A., Paulovich, A., Pomeroy, S.L., Golub, T.R., Lander, E.S., et al. (2005). Gene set enrichment analysis: A knowledge-based approach for interpreting genome-wide expression profiles. Proc. Natl. Acad. Sci. U. S. A. 102, 15545–15550.

