## Supplementary_figures for "SUMO2 Inhibition Reverses Aberrant Epigenetic Rewiring Driven by Synovial Sarcoma Fusion Oncoproteins and Impairs Sarcomagenesis"

S1

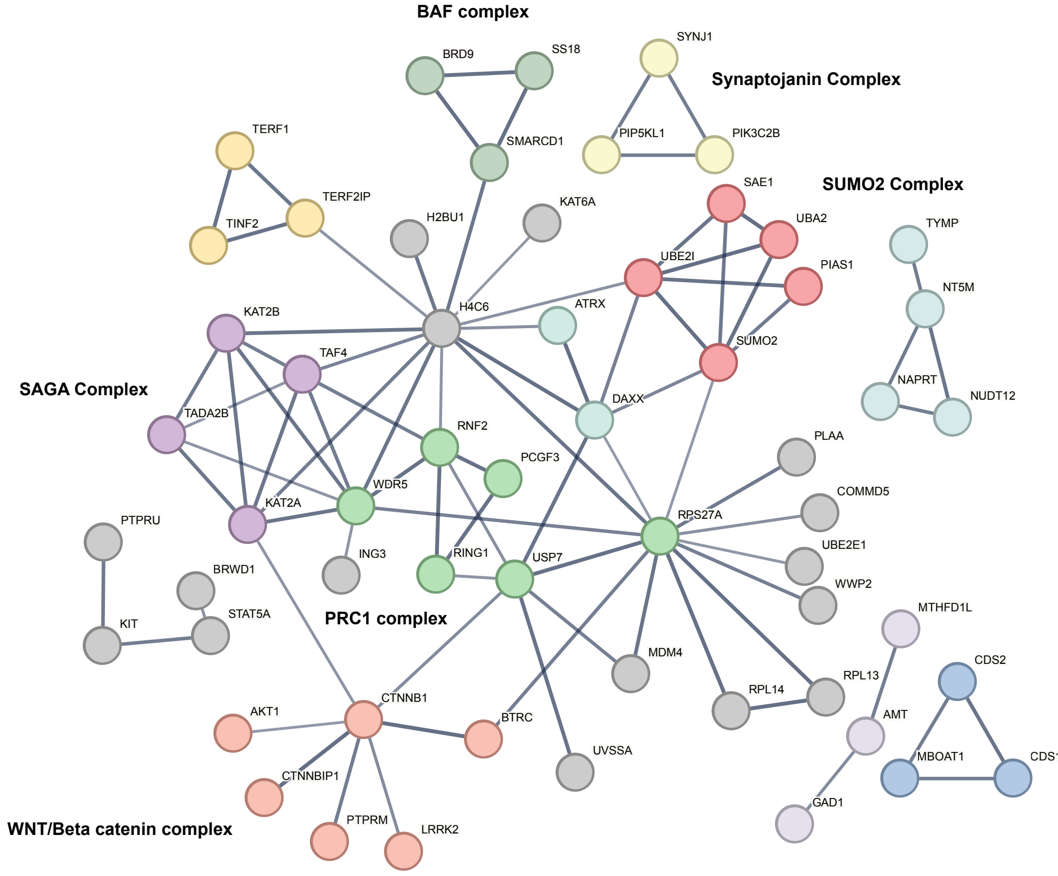
BAF complex
BRD9
SS18
SMARCD1
SYNJ1
Synaptojanin Complex
PIP5KL1
PIK3C2B
TERF1
TERF2IP
TINF2
H2BU1
KAT6A
SAE1
SUMO2 Complex
UBA2
UBE2I
PIAS1
TYMP
NT5M
KAT2B
H4C6
ATRX
SUMO2
TAF4
SAGA Complex
TADA2B
DAXX
NAPRT
NUDT12
RNF2
PCGF3
PLAA
WDR5
COMMD5
KAT2A
RPS27A
PTPRU
UBE2E1
ING3
RING1
USP7
WWP2
BRWD1
MTHFD1L
STAT5A
PRC1 complex
KIT
MDM4
CDS2
RPL13
RPL14
AMT
AKT1
CTNNB1
BTRC
MBOAT1
CDS1
UVSSA
CTNNBIP1
GAD1
PTPRM
LRRK2
WNT/Beta catenin complex

**S2**

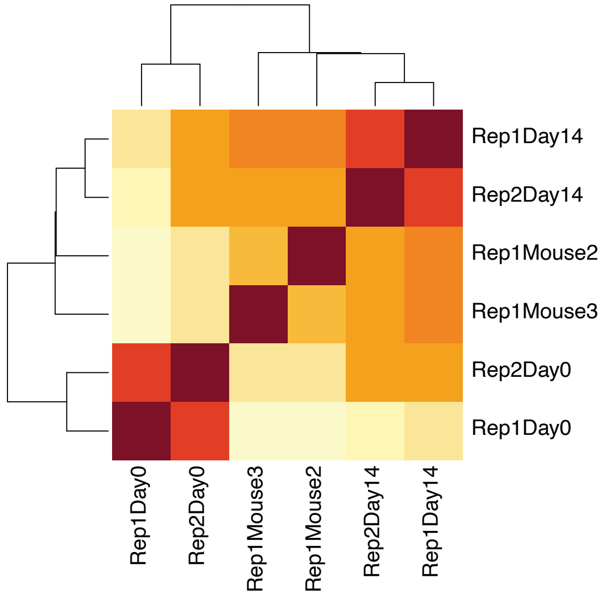
Rep1Day14
Rep2Day14
Rep1Mouse2
Rep1Mouse3
Rep2Day0
Rep1Day0
Rep1Day0
Rep2Day0
Rep1Mouse3
Rep1Mouse2
Rep2Day14
Rep1Day14

**S3**

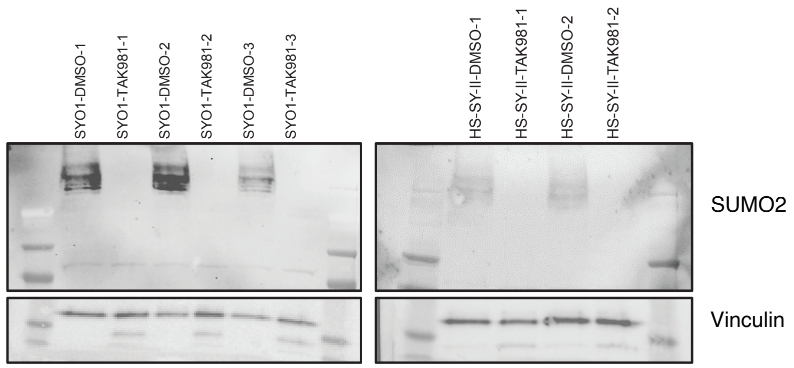
SYO1-DMSO-1
SYO1-TAK981-1
SYO1-DMSO-2
SYO1-TAK981-2
SYO1-DMSO-3
SYO1-TAK981-3
HS-SY-II-DMSO-1
HS-SY-II-TAK981-1
HS-SY-II-DMSO-2
HS-SY-II-TAK981-2
SUMO2
Vinculin

**S4**

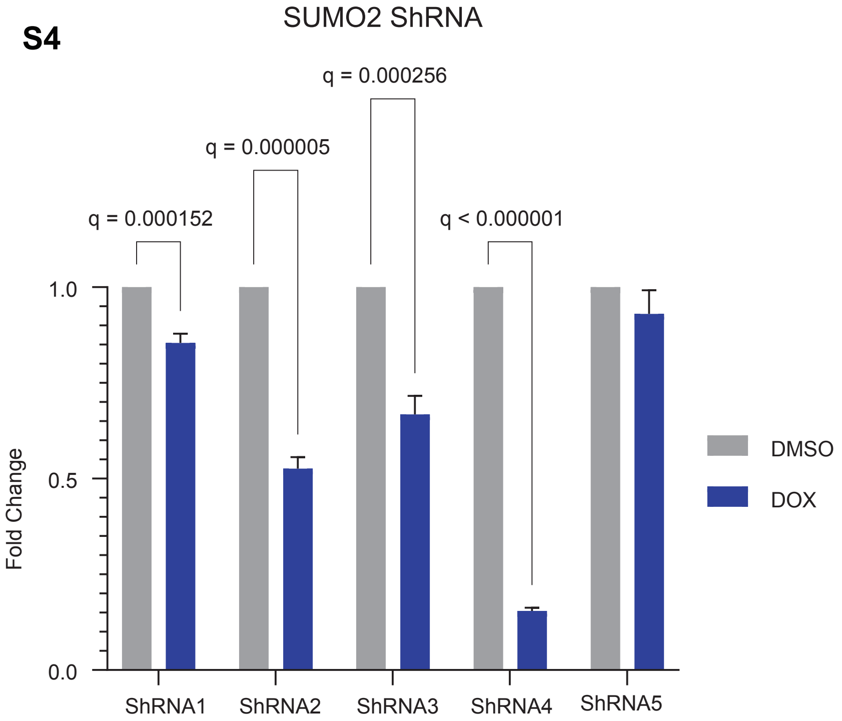
SUMO2 ShRNA
q = 0.000152
q = 0.000005
q = 0.000256
q < 0.000001
Fold Change
1.0
0.5
0.0
ShRNA1
ShRNA2
ShRNA3
ShRNA4
ShRNA5
DMSO
DOX

**S5**

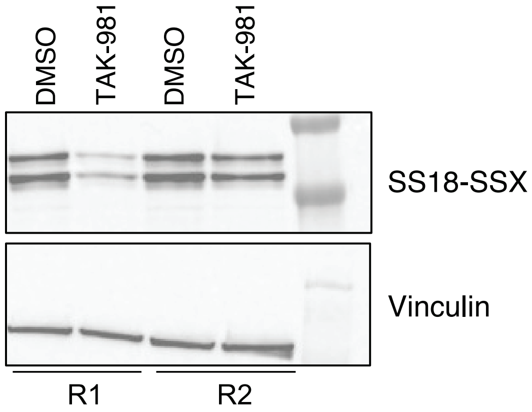
DMSO
TAK-981
DMSO
TAK-981
SS18-SSX
Vinculin
R1
R2

**S6**

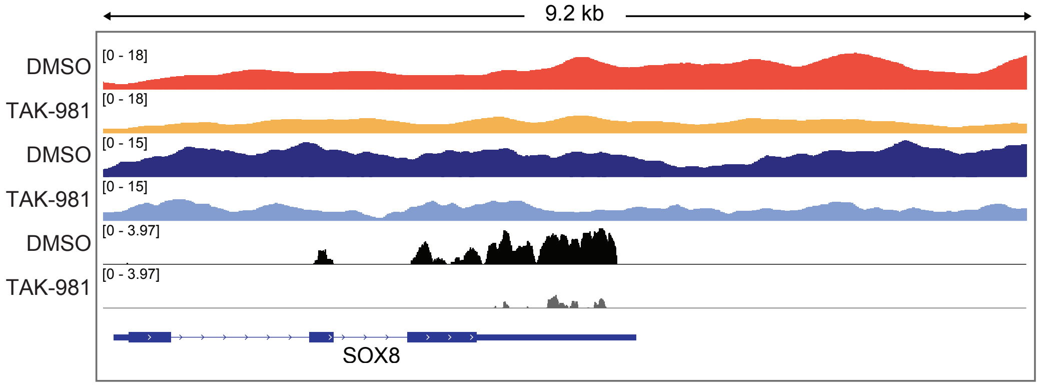
9.2 kb
DMSO
[0 - 18]
TAK-981
[0 - 18]
DMSO
[0 - 15]
TAK-981
[0 - 15]
DMSO
[0 - 3.97]
TAK-981
[0 - 3.97]
SOX8

**S7**

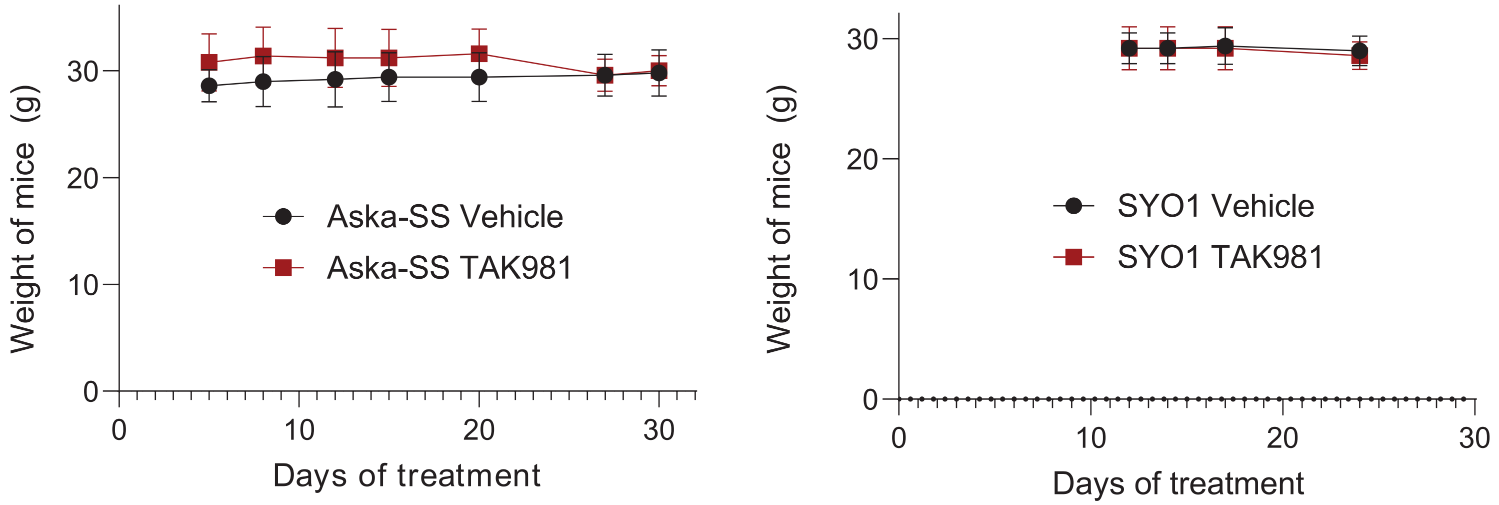
Weight of mice (g)
30
20
10
0
Aska-SS Vehicle
Aska-SS TAK981
Days of treatment
Weight of mice (g)
SYO1 Vehicle
SYO1 TAK981
Days of treatment

### Supplemental Figure Legends:

**Figure S1:**

String analysis of synovial sarcoma selective genes identified in our study is shown. Nodes indicate genes, edges indicate connections between the genes as represented by participation in common protein complexes identified in deposited experimental data from other groups.

**Figure S2:** Correlation matrix of replicates from the *in vitro* and *in vivo* screen is shown. Rep = replicate. Day 0 = start of the assay and D14 is the end of the *in vitro* assay.

**Figure S3:** Immunoblotting for total SUMO2 in lysates from SYO1 cells treated with TAK-981 compared to DMSO control are shown with vinculin as the loading control.

**Figure S4:** Fold changes (Y-axis) of SUMO2 transcripts in doxycycline-treated relative to DMSO-treated HS-SY-II cells as measured by quantitative PCR are shown with 5 independent SUMO2-targeting shRNAs is shown (X-axis). Doxycycline was used to induce the shRNA from a tetracycline-responsive promoter.

**Figure S5:** Immunoblot analysis of whole-cell lysates from 1273/99 cells treated with TAK-981 and probed for the SS18-SSX2 fusion protein are shown. Vinculin is shown as a loading control.

**Figure S6:** Integrated genome viewer (IGV) tracks for the SS18-SSX fusion and H2AK119ub in DMSO or TAK-981 treated SYO1 cells along with corresponding RNAseq tracks are shown for SOX8, a SS18-SSX target gene

**Figure S7:** Weights of mice (grams, Y-axis), treated with TAK-981 (scarlet) compared to DMSO (black) lines are shown at various time intervals (days, X-axis).
